# Transiently increased intercommunity regulation characterizes concerted cell phenotypic transition

**DOI:** 10.1101/2021.09.21.461257

**Authors:** Weikang Wang, Ke Ni, Dante Poe, Jianhua Xing

**Affiliations:** Department of Computational and Systems Biology, University of Pittsburgh, Pittsburgh, PA 15232, USA; CAS Key Laboratory of Theoretical Physics, Institute of Theoretical Physics, Chinese Academy of Sciences, Beijing 100190, China; Joint CMU-Pitt Ph.D. Program in Computational Biology, University of Pittsburgh, Pittsburgh, PA, USA; Department of Physics and Astronomy, University of Pittsburgh, Pittsburgh, PA 15232, USA; UPMC-Hillman Cancer Center, University of Pittsburgh, Pittsburgh, PA, USA

## Abstract

Phenotype transition takes place in many biological processes such as differentiation and reprogramming. A fundamental question is how cells coordinate switching of expressions of clusters of genes. Through analyzing single cell RNA sequencing data in the framework of transition path theory, we studied how such a genome-wide expression program switching proceeds in five different cell transition processes. For each process we reconstructed a reaction coordinate describing the transition progression, and inferred the gene regulation network (GRN) along the reaction coordinate. In all processes we observed common pattern that the overall effective number and strength of regulation between different communities increase first and then decrease. The change accompanies with similar change of the GRN frustration, defined as overall conflict between the regulation received by genes and their expression states, and GRN heterogeneity. While studies suggest that biological networks are modularized to contain perturbation effects locally, our analyses reveal a general principle that during a cell phenotypic transition, intercommunity interactions increase to concertedly coordinate global gene expression reprogramming, and canalize to specific cell phenotype as Waddington visioned.

## INTRODUCTION

A lasting topic in science and engineering is how a dynamical system transits from one stable attractor to a new one in a corresponding state space [1]. For example, a substitution reaction in organic chemistry can proceed either through first breaking an existing chemical bond to form an intermediate planar structure followed by forming a new bond, termed as a SN1 mechanism, or through forming a trigonal bipyramidal intermediate complex where breaking of the old bond and formation of the new bond take place concertedly, termed as a SN2 mechanism (Fig. 1a) [2]. Which mechanism dominates a process is determined by both the relative thermodynamic stability of the two intermediate structures, and the kinetics of forming them.

**Figure 1.**
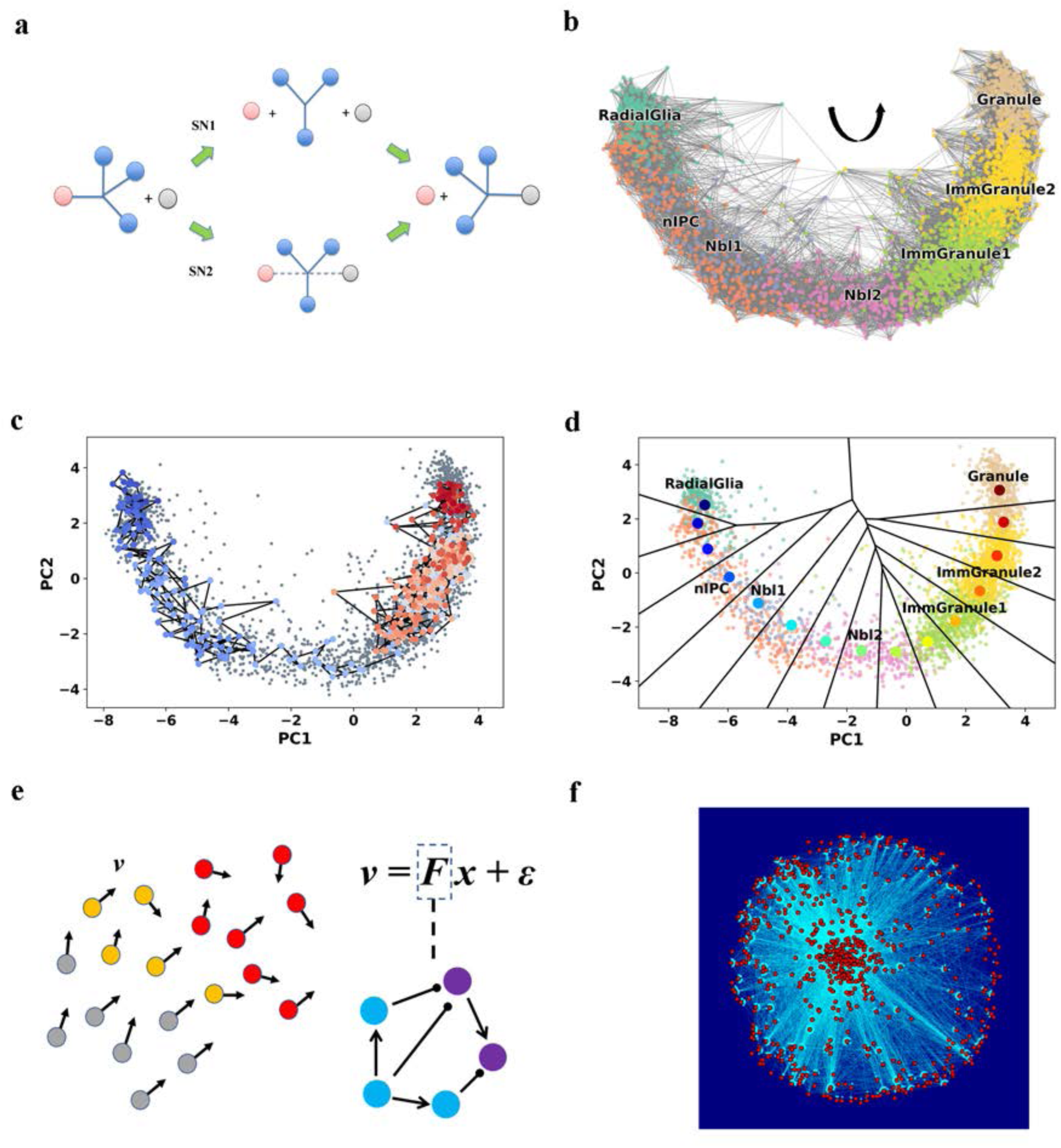
Dynamical analyses of scRNA-seq data of dentate gyrus neurogenesis. (a) Competing SN1 and SN2 mechanisms for substitution reactions. The nodes and edges represent chemical groups and chemical bonds respectively. (b) scRNA-seq data and RNA velocity-based transition graph shown in the cell expression state space (shown in the 2D leading principal components (PCs) space). Each dot represents a cell, and each edge between two dots indicates a transition between cell states corresponding to the two cells. Color represents cell type. Arrow indicates direction of transition. (c) A typical single cell trajectory simulated on the transition graph, illustrated in the 2D leading PCs space. The trajectory starts from the Radial-Glia cell type and transits into the granule cell type. The dot color represents the progression of the trajectory (from blue to red). (d) 1-D RC reconstructed from the simulated single cell trajectories by using a revised finite temperature string method. The large colored dots represent the RC points (start from blue and ends in red). Voronoi grids are generated with the RCs. The small dots are cells (color represent cell type). (e) Schematic plot of GRN inference. The colored dots with arrow are cells with their RNA velocities (different colors represent different cell types). (f) Plot of inferred GRN of dentate gyrus neurogenesis. Each red dot represents a gene and each line stands for regulation between two genes.

Another examples of transitions that attracts increased interest recently are transitions between different cell phenotypes, partly due to available genome-wide characterization of the cell gene expression state throughout a transition process aided with advances of single cell genomics techniques [3–5]. A cell is a nonlinear dynamical system governed by a complex regulatory network. The latter is formed by a large number of interacting genes, and can have multiple stable attractors corresponding to different cell phenotypes [6, 7]. Typically a large number of phenotype-specific genes maintain a specific phenotype through mutual activation while suppressing expression of genes corresponding to other exclusive phenotypes [8]. In some sense it resembles a spin system segregating into upward and downward domains. When a cell phenotypic transition (CPT) takes place, the genes need to switch their expression status, analogous to flipping some upward and downward spin domains [9, 10].

A question arises as how a CPT, or a cell state transition process in general, proceeds. Answering this question requires examining how a genome-wide gene regulatory network (GRN) changes during a CPT. The transition may be sequential with deactivation of regulatory network links between genes in pairs of functional modules, followed by activation of network links between other pairs of functional modules to instruct the cell into one specific meta-stable state. For example, it has been noticed that cell fate decision making preferentially takes place in the G1 phase with recruitment of transcription factors (TFs) that rely on CDK (cyclin-dependent protein kinase) to activate genes of the new cell type, following the M phase where TFs disassociate from the condensed chromatin allowing reset of the cell type specific expression program at the transcription level [11]. Alternatively, deactivation and activation of regulatory network links may happen concurrently as in the SN2 mechanism [5, 8, 12]. One can vision two qualitatively different characteristics of the two mechanisms.

Developments of single cell RNA sequencing (scRNA-seq), together with analysis tools, have expanded our knowledge on CPT [13–16]. Most of these tools, such as Monocle, Scanpy and Seurat, focus on the pseudo-time analysis given that the scRNA-seq data only provides snap-shot information [3, 13, 15, 17–22]. In this work we exploited a recently developed RNA velocity formalism [23]. The formalism utilizes the number of un-spliced and spliced message RNAs to infer the time derivative of a cell state in the gene expression space, and predicts the single cell state in the next few hours. RNA velocity has been used for prediction of cell state and reveals information of dynamics of CPTs such as differentiation [24, 25]. Using the RNA velocity data we further developed a dynamical model, and analyzed the transition dynamics in the framework of reaction rate theories and network science theories. A concerted mechanism is supported by characterizations of five CPT processes with a number of statistical quantities, notably a conserved pattern of peaked intercommunity interactions at an intermediate stage of each transition.

## RESULTS

### Dynamical model reconstructed from scRNA-seq data describes development of granule lineage in dentate gyrus

Dentate gyrus locates in the brain hippocampus, and is important for memory formation [26, 27]. During the process of neurogenesis, radial glia-like cells differentiate through nIPCs, neuroblast 1 and 2, immature granule cells, and eventually into mature granule cells (Fig. 1b) [28]. A number of transcription factors such as Hes5, Sox2 and CREB play important roles at various steps of this process [29].

First, we constructed a dynamical model from the dataset of Hochgerner et al. [28]. Using the package dynamo[30], we constructed a transition graph *T* through RNA velocity analyses, and modeled the developmental processes and Markovian transitions among the sampled cells [25, 31] (Fig. 1b). We performed stochastic simulations on this transition graph and obtained an ensemble of transition trajectories (Fig. 1c, Materials and Methods). Next, we used reaction coordinate (RC, denoted by {*r*}, where *r* is the index of the discretized RC) to characterize the progression of granule lineage development. RC is a central concept in rate theories [1] that uses a one-dimensional manifold to reflect progression of the reaction process, and shares some similarity to the pseudo-time trajectory concept in the single cell genomics field [13, 15]. We calculated an array of RC points from the simulated trajectories, which reflects the continuous progression from Radial Glia to Granule cells through a set of intermediate cell types (Fig. 1d) [32–34].

Next, we set to further analyze how the regulatory network reorganizes during this process. Qiu et al. showed that one can extract quantitative information about gene regulation from single cell expression and RNA velocity data, which agree well with available experimental studies when tested on several datasets [30]. We adopted a model simpler than that of Qiu et al.[30] by assuming the governing equation as ***v*** = ***Fx*** + ***ε*** , with ***F*** for the intracellular gene regulatory network with each element *F_ij_* quantifying the regulation direction and strength of gene *j* on gene *i* in the GRN, and *ɛ* being random white noises. We obtained ***F*** through partial least square regression (PLSR) [35] over the scRNA-seq data and estimated RNA velocities (Materials and Methods) (Fig. 1e). Figure 1f shows the inferred genomewide GRN of this process, and some of the inferred regulations have literature support (Table S1).

### Network analyses reveal increase of frustration and network heterogeneity during transition

Similar to the SN1 v.s. SN2 mechanisms, the concerted but not the sequential mechanism predicts significantly increased gene-gene interactions. Therefore, to evaluate the two possible mechanisms, we binarized gene expression (0 for silence, and 1 for active expression, Fig. 2a) (denoted as s*_i_* for gene *i*) and calculated the number of effective edges, i.e., edges with nonzero *F_ij_* and gene *j* expressed (*s_j_* = 1) in the cell, which indeed increases first and then decreases along the RCs (Fig. S1a).

**Figure 2.**
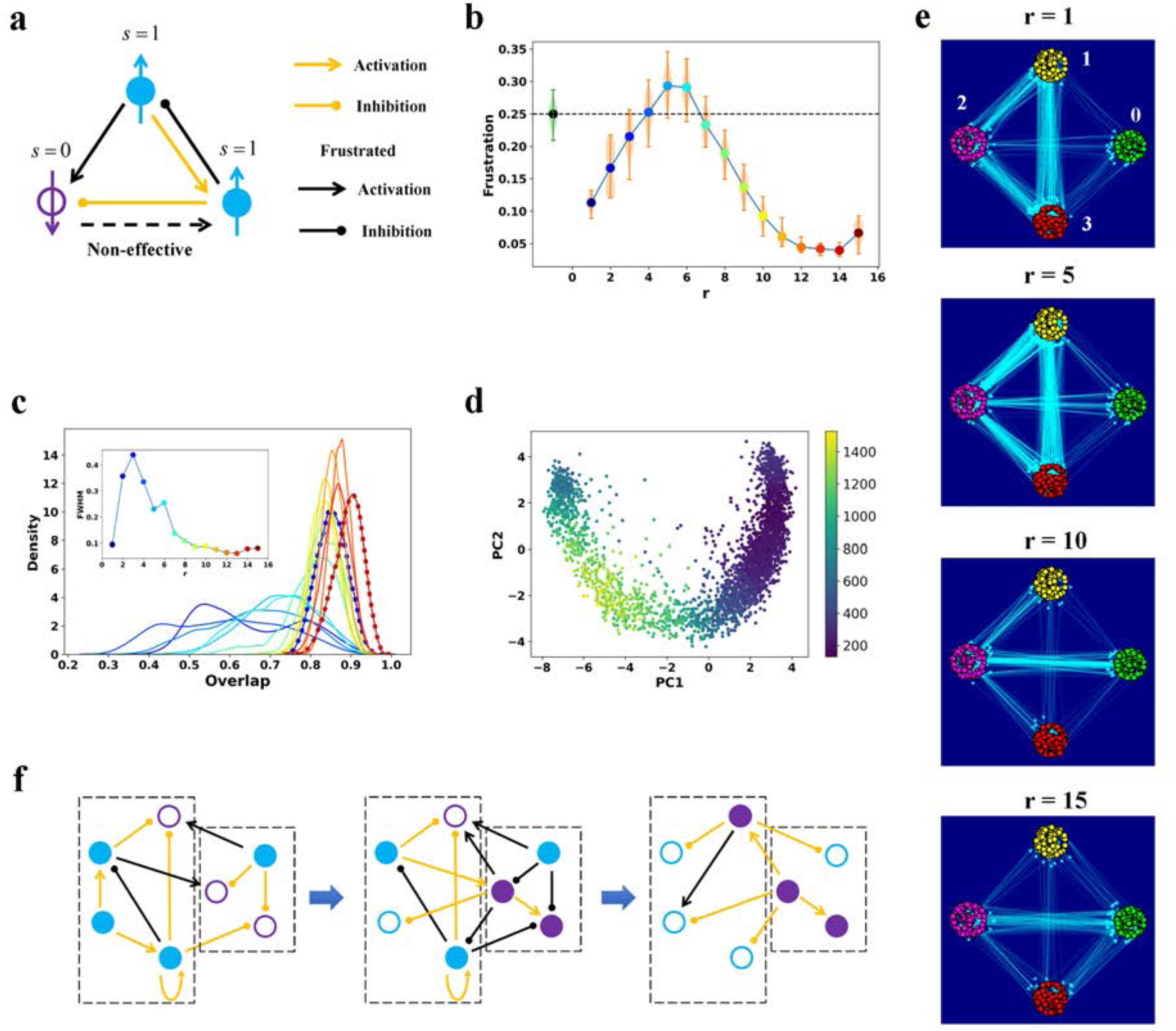
Analyses of scRNA-seq data of dentate gyrus neurogenesis reveal a concerted transition mechanism. (a) Schematic plot of frustration in GRN. Filled circles (Upwards arrow) represent active genes. Empty circle (Downwards arrow) represents a silent gene. Colors represent marker genes of different cell states (light-blue circle represent genes are active in initial states and purple circles represent genes are active in final state). (b) Frustration score along the RC of dentate gyrus neurogenesis. The mean and variance at each RC point were calculated using k-nearest-neighboring cells of the corresponding RC point within the Voronoi grid. Green violin represents distribution and dashed line is the average frustration score of random samples. (c) Variation of distribution of state overlap along the RC of dentate gyrus neurogenesis. Colors of the distributions correspond to that of RC points. The initial and final RCs are plotted with dotted line. Inserted graph is the FWHM of the distributions along the RC. (d) Cell specific variation of the number of effective intercommunity edges (represented with color) in the GRN of each single cell. (e) Evolution of the number of effective intercommunity edges along the RC during dentate gyrus neurogenesis. Each node represents a gene. Color of the node represents index of community. Arrow represents direction of regulation. r is the index in the RC. (f) Schematic of the concerted mechanism for a cell phenotypic transition. The dash-line boxes represent communities. Legends are the sample as Fig. 2a.

In the SN2 mechanism, the increased bond number is due to coexistence of the bonds that are to form and break. Analogously, for a concerted mechanism one expects co-expression of genes that normally only express in one stable phenotype, leading to conflict on the expression state of a gene and the regulation acting on it. We used frustration to quantify such conflicting regulations (Materials and Methods). That is, regulation from gene *j* to gene *i* is frustrated if the expression state of gene *i* ( *s_i_* ) contradicts the regulation from gene *j* [36]. For instance, if gene *i* is active but regulation from an active gene *j* (with *s_j_* = 1) is inhibition, this regulation is frustrated.

Furthermore we defined the overall frustration score of a cell-specific GRN as the fraction of frustrated edges out of all edges in the whole network of the cell [36]. For development of the granule lineage, the average frustration score along RCs increases first and reaches a peak corresponding to neuroblast cells, then decreases (Fig. 2b), consistent with the concerted but not the sequential mechanism. For comparison we also calculated the frustration score of random cell states (green violin plot and the dashed line in Fig. 2b). Notice that some of random states may not be biologically accessible. The frustration score of the cellular system is lower in the initial and final metastable states compared to that of the random states, but reaches a peak value higher during the transition.

During transition, exploration of state space by cells leads to increased cell state heterogeneity in this process [8]. We used a distribution of state overlap to quantify the heterogeneity during CPT. Originally used in spin glass and Boolean state model to characterize the state structures [9, 37], the state overlap is defined as 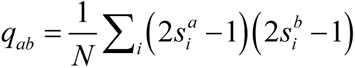, where *a* and *b* represent different cells sampled in the Voronoi grid of each RC point, *N* is the number of genes, and the sum is over all the genes included in the analyses. To be consistent with the definition in spin glass, 2*s_i_* −1 (value equals 1 or -1) is used in calculation instead of *s_i_* q*_ab_*. equals 1 if the cell *a* and *b* are indentical, and q*_ab_* equals -1 if the state vectors of cell *a* and *b* are the opposite. A lower heterogeneity gives a narrower distribution and a higher mean value of the state overlap. Figure 2c shows the distribution of state overlap P*_r_* (q*_ab_* )calculated at each RC point. During transition, first the distribution along the RC becomes more dispersive and its mean value approaches to 0 from 1 gradually. Then the dispersion narrows down and returns to a mean value close to 1. This trend is apparent from the variation of full width at half maximum (FWHM) of the P*_r_* (q*_ab_* ) along RC (Fig. 2c inserted). This reflects a temporal increase of heterogeneity along the transition. We also quantified GRN network heterogeneity [38, 39]. Network heterogeneity measures how homogenously the connections are distributed among the genes [38]. A high heterogeneity indicates coexistence of hub genes with high connectivity and genes with low connectivity. The network heterogeneity increases first, reaching a maximum neuroblast cells, then decreases when approaching the mature granule cell state (Fig. S1b) [38]. We then observed a similar pattern using another type of heterogeneity called degree heterogeneity, for which a higher value reflects a more uneven distribution of the degree of genes in a GRN, and the degree of a gene is the number of connections this gene has to other genes (Fig. S1c)[39]. This result further indicates that the trend is not due to a specific choice of the heterogeneity.

### Reconstructed gene regulatory network reveals increased intercommunity interactions at intermediate stage of transition

To examine the nature of the increased interactions, we divided the inferred GRN into four communities using the Leiden method (Materials and Methods) [40, 41]. The number of effective intra-community edges correlates with the number of active genes (Fig. S1d), but we did not observe a universal pattern among the four communities on how intracommunity interactions change along the RCs. Actually, the number of intracommunity edges for community 0 increases, that of community 2 decreases, while those of the community 1 and 3 peak in the middle (Fig. S1d left). In contrast, the number of effective inter-community edges increases first and then decreases between pairs of all four communities (Fig. 2d). The intercommunity interaction strengths, defined as the total number of effective edges between different communities, are the strongest at *r = 5* (Fig. 2e). This variation of intercommunity effective edges does not correlate with the total number of active genes, with a correlation coefficient only 0.26 (Fig. S1e).

To rule out the possibility that the observed properties are specific for the Markov model we used, we repeated the above analyses with a transition matrix reconstructed from a correlation kernel of Bergen et al. [42]. The analyses gave the RCs, and patterns of the frustration score and the number of effective inter-community edges changes along the RC similarly as what observed with the model obtained using that of Qiu et al. (Fig. S2a-c).

Figure 2f shows a schematic summary of the concerted mechanism. Co-expression of conflicting genes leads to increased intercommunity edges, and frustrated edges. Some of the genes transiently act as hub genes, which lead to increased network heterogeneity.

### Reconstructed gene regulatory network reveals similar concerted mechanism in other CPT systems

To investigate whether the concerted mechanism is general for CPTs, we performed the same analyses on a number of additional CPT scRNA-seq datasets. One system is hematopoiesis. Hematopoietic stem cells (HSCs) mainly locate in the bone marrow and can differentiate into different types of blood cells including the erythrocytes and lymphocytes[43]. In this process, the lineage choice is controlled by multiple signals such as Notch and IL2, and regulated by different transcriptional factors such as GATA2 and E2A [44, 45]. In this dataset, we analyzed the transition from HSCs to monocytes via precursors (Fig. 3a) [46]. Along the RC calculated from the data (Fig. 3b), the cells show the highest frustrations score when they almost exit the stage of precursor (Fig. 3c). The dispersion of P*_r_* (q*_ab_* ) also reaches a maximum value during the course of hematopoiesis (Fig. 3d). The number of effective inter-community edges increases first and then decreases (Fig. 3e&3f). The number of effective edges also shows similar trend (Fig. S3a). Furthermore, the progression of network heterogeneity along the RC behaves similarly (Fig. S3b&3c).

**Figure 3.**
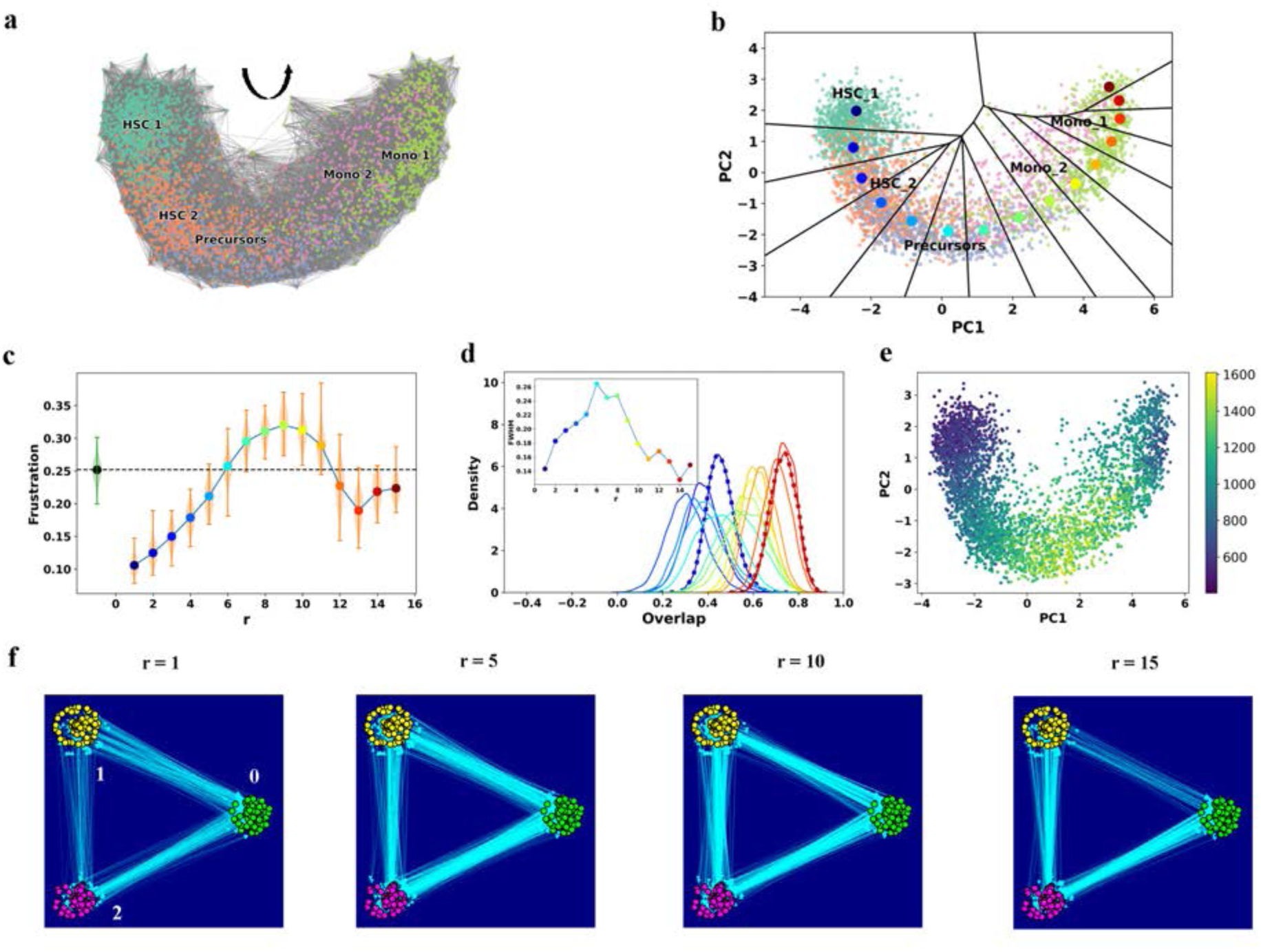
Analyses on hematopoiesis dataset reveals features of a concerted mechanism. (a) Transition graph of hematopoiesis based on RNA velocity. Color represents cell type. Arrow indicates the direction of transition. (b) RC (large colored dots, start from blue and ends in red) of hematopoiesis with corresponding Voronoi grids. Small dots are cells (color represents cell type). (c) Frustration score along the RC in hematopoiesis. (d) Variation of distribution of state overlap along the RC in hematopoiesis. Colors of the distributions correspond to that of RC points. The initial and final RCs are plotted with dotted line. Inserted graph is the FWHM of the distributions along the RC. (e) Cell-specific variation of effective intercommunity regulation in hematopoiesis. Color represents the number of effective intercommunity edges within each cell in the GRN. (f) Evolution of the number of effective intercommunity edges along the RC during hematopoiesis. Each node represents a gene. Color of the node represents index of community. Arrow represents direction of regulation. r is the index in the RC.

To confirm the results, we tested another GRN inference method GRISLI [47]. For the datasets tested, the trend that frustration score and the inter-community edges increase first and decrease remains the same as that with PLSR (Fig. S4a-b).

The datasets above are from mouse samples. We also performed the same analyses on development of human glutamatergic neurogenesis. The dynamics of inter-community edge number and frustration score along the RC shows similar trends (Fig. S5a-d). We also analyzed an induced in-vitro transition process, the Epithelial-to-Mesenchymal Transition (EMT) of human A549 cells treated with TGF-β for different durations [48]. This dataset composes a total of *N = 3003* single cell samples measured at several time points (Fig. S6a). EMT takes place in a number of biological and pathological processes. The process is regulated by some central transcription factors including Snail, Zeb, and Twist, and proteins like E-cadherin and vimentin have been identified as important markers for the epithelial and mesenchymal phenotypes, respectively [49]. Studies in recent years demonstrate existence of intermediate states called partial EMT states besides the two terminal epithelial and mesenchymal phenotypes during the process of EMT[49–53]. During EMT, cells change from epithelial to mesenchymal phenotypes with an increased EMT hallmark gene set score. An array of RC points connects the epithelial (cells without treatment or 0 day) and mesenchymal states (cells treated with TGF-β for 3 days) (Fig. S6b). Along the calculated RC, the frustration score shows a peak during transition (Fig. S6c). The number of effective inter-community edges increases first and then decreases when epithelial cells transit into mesenchymal cells (Fig. S6d). Compared to results from other datasets, all quantities calculated from this dataset show larger variances, probably due to the larger expression heterogeneity in cultured cancer cells.

### Pancreatic endocrinogenesis shows mixed features of concerted and sequential mechanisms

Another dataset we analyzed is the development of pancreatic endocrine cells (Fig. 4a) [54]. Neurogenin3 (Ngn3) is a transcription factor that transiently expresses in endocrine progenitor-precursor [55]. Transcription factor Fev is reported as a marker of the intermediate state between initial endocrine progenitors and the final state such as differentiated α-cells and β-cells [55, 56]. As characterized by the RC (Fig. 4b), during embryonic development Ngn3-low progenitors first transform into Ngn3-high precursors then Fev-high (Fev+) cells. The latter further develop into endocrine cells including glucagon producing α-cells and β-cells.

**Figure 4.**
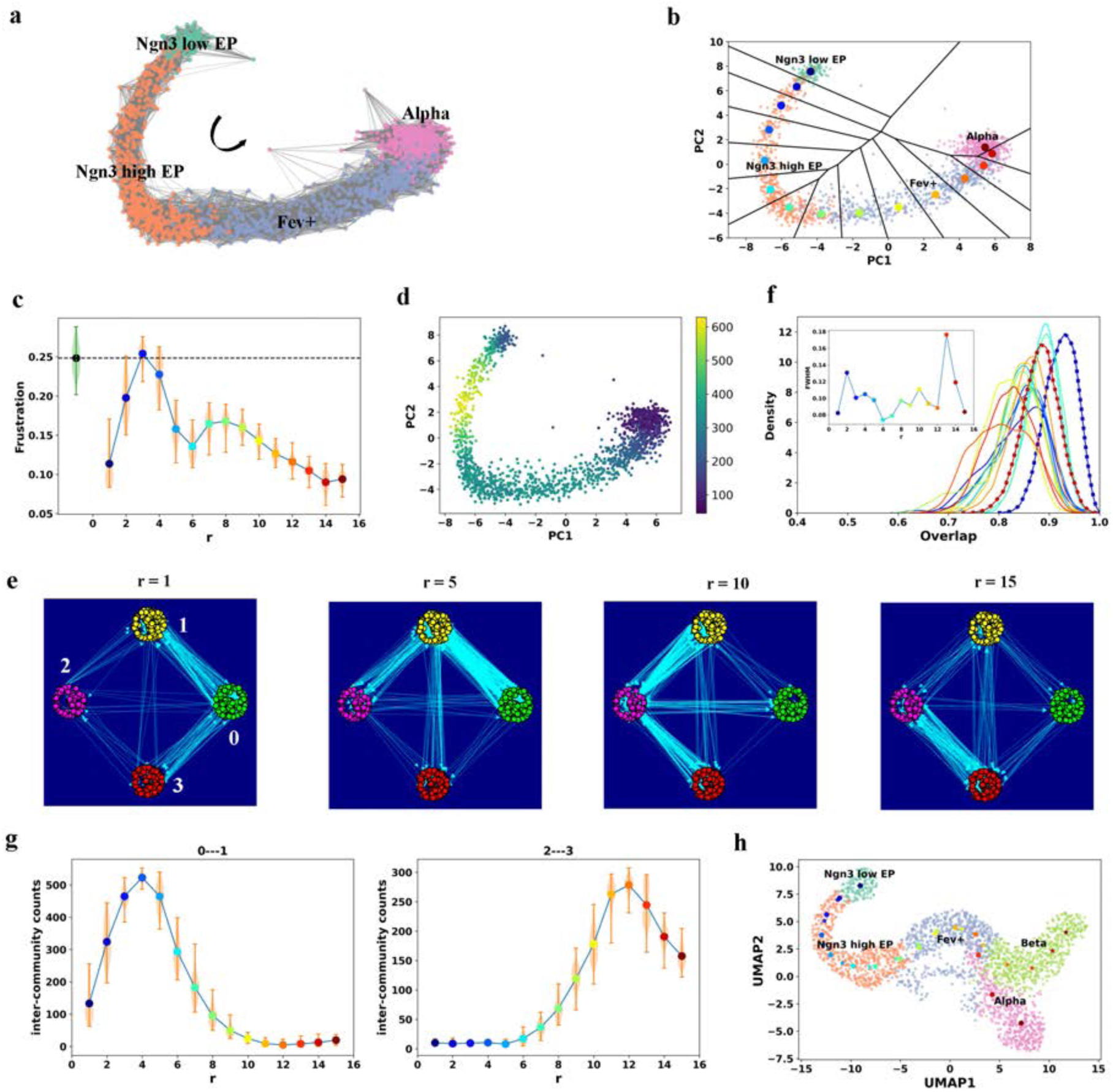
The scRNA-seq dataset of pancreatic endocrinogenesis reveals features of mixed concerted and sequential mechanisms. (a) Transition graph of pancreatic endocrinogenesis based on RNA velocity. Color represents cell type. Arrow indicates the direction of transition. (b) RC (large colored dots, start from blue and ends in red) of pancreatic endocrinogenesis with corresponding Voronoi grid. Small dots are cells (color represents cell type). (c) Frustration score along the RC in pancreatic endocrinogenesis. (d) Cell-specific variation of effective intercommunity regulation in endocrine cell development. Color represents the number of effective intercommunity edges within each cell in the GRN. (e) Evolution of the number of effective intercommunity edges along the RC during pancreatic endocrinogenesis. Each node represents a gene. Color of the node represents index of community. Arrow represents direction of regulation. r is the index in the RC. (f) Variation of distribution of state overlap along the RC in pancreatic endocrinogenesis. Colors of the distributions correspond to that of RCs. The initial and final RCs are plotted with dotted line. Inserted graph is the FWHM of the distributions along the RC. (g) Number of effective intercommunity edges between community 0 and 1 (left) and that between community 2 and 3 (right) along the RC during pancreatic endocrinogenesis. (h) RCs of both α (large round dots) and β (pentagram dots) branch of pancreatic endocrinogenesis projected on embedding plane of UMAP (Uniform Manifold Approximation and Projection). Small dots are cells and color represents cell type.

At a first glance, the frustration score and overall inter-community interactions show similar trend as other CPT systems analyzed previously (Fig. 4c). The number of effective inter-community edges increases first and then decreases while low Ngn3 expression cells transit into α-cells (Fig. 4d-e). The number of effective edges and the network heterogeneity show similar trends (Fig. S7a-c). However, a closer examination reveals a new pattern. Even though the variance of the violin plot is large, the average frustration score reaches a local minimum at *r = 6* corresponding to the cell population with high Ngn3 expression. Results with the GRN inferred with GRISLI also show a similar trend (Fig. S8a). Furthermore, the P*_r_* (q*_αβ_* ) along the RCs not only has low dispersion values at the initial and final states, but also exhibits a local minimum at *r = 6* (Fig. 4f), which is different from other datasets.

Notice for the initial state (*r = 1*), strong interactions exist between community 0 and community 1 (0-1 interactions), and medium interactions exist between community 0 and 3 (0-3 interactions). Compared to the initial state, the final state (*r = 15*) shows decreased 0-1 and 0-3 interactions, and increased 2-3 interactions. Along the RC, we do not observe coexistence of these interactions expected for a concerted mechanism (Fig. 4g and Movie S1). Instead, the transition is mediated through transient increase of 0-2 and 1-2 interactions. The 2-3 interaction shows significant increase only when *r* exceeds 6, while the 0-1 interaction already decreases and falls to a relatively low value (Fig. 4g). Analyses on GRN inferred with GRISLI reveal similar results (Fig. S8a-c). Clearly, the pancreatic endocrinogenesis and other datasets analyzed above show qualitatively different patterns of intercommunity interactions along the RC. On the other hand, one may consider the overall transition as a sequential of two transitions, i.e., Ngn3-low progenitors first becomes a “primed” Ngn3-high state, then the latter differentiates into α-cells. For each transition the intercommunity interactions and the frustration score increase transiently.

The results in Fig. 4 b-g lead to a hypothesis that the primed Ngn3-high state may act as a common initial state that can differentiate into multiple terminal cell types, analogous to the intermediate of the SN1 mechanism. To examine this hypothesis, we analyzed the branch that differentiated into glucagon producing β-cells (Fig. S9a-d). Here again the genes are grouped into four communities. The genes in community 0, 1 and 2 have large proportion overlap with that in the α branch, respectively, consistent with the observation that the two branches share the same “trunk” including Ngn3-low progenitors, Ngn3-high precursors, and Fev+ cells. The genes in community 3 of the β branch mainly show high expression in the glucagon producing β-cells, which differ from those in community 3 of the α branch mainly showing high expression in the glucagon producing α-cells (Fig. S9e). In this β branch, we also observed similar sequential variation on the frustration score and inter-community edges. The effective 2-3 interactions start to increase when 0-1 interactions decrease (Fig. S9f). These results show that genes associated with different cell states are suppressed before cells approach to the “cross” (Fig. 4h, for better visualization, we use UAMP embedding here).

## DISCUSSIONS

The idea of relating CPTs and chemical reactions has been discussed in the literature [8]. Here we presented a procedure of reconstructing the RC of a CPT process from scRNA-seq data. A related concept is the transition state. In chemical reactions it typically refers to short-lived intermediates or a state of maximal potential energy along the RC. It is tempting to identify the intermediate state with highest frustration as the “transition state”, while it is unclear whether it is indeed a dynamical bottleneck of the associated CPT process. Compared to a chemical reaction, a CPT process is more complex. Either the concerted or the sequential mechanism may be extreme cases, and some CPTs may involve gradual changes of the expression program. With the quantitative analysis pipeline developed in this work one may systematically enumerate the transition patterns. Our study shows that collective quantities such as the frustration score characterize CPT dynamics, similar to the identification of single cell transcriptional diversity as a hallmark of developmental potential [57].

In a series of studies, geometric analyses on the differential equations governing a CPT also identify two basic structures [58, 59]. The binary choice mechanism is a direct conversion between two states through an intermediate saddle point, corresponding to the concerted mechanism discussed in this work. The binary flip mechanism describes a transition route emanating from the original state to an intermediate then two possible final states, sharing some basic features we observed for the sequential transitions in pancreatic endocrinogenesis. Therefore, two different analyses focusing on different aspects of CPTs have reached similar conclusions on the archetypal designs for cell fate decisions. The present work focuses on how the gene expression pattern changes during a CPT process, without reference on the nature of the intermediate transition states, i.e., whether they are metastable attractors or saddle points. Due to the quality of the scRNA-seq data, we could not use dynamo to reach a decisive conclusion on the identities of these intermediates and geometric structure of the vector field of the dynamical systems corresponding to the datasets we analyzed here.

The observed common pattern of transiently peaked intercommunity interactions provides a new angle to examine the structure-function relation of a biological network. Previous theoretical and experimental studies have shown that a biological network is generally modularized with dense intracommunity interactions and sparse intercommunity interactions, which helps insulating perturbations in one community from propagating globally and increases functional robustness of each module [60–62]. The observed pattern supports that the number of intercommunity interactions of a GRN is smaller at stable phenotypes than that of intermediate transient phenotypes.

During a CPT, a cell needs to escape a stable phenotype, and the increased intercommunity interactions help on coordinating gene expression profile change among communities. The decreased modularity is consistent with a critical state transition mechanism [7] that individual components become more connected and correlated, as what observed near the critical point of a phase transition. Even though different CPTs mainly show concerted variation, some CPTs such as pancreatic endocrinogenesis may adopt a mixed strategy of concerted and sequential mechanisms. Further studies are needed to investigate how the coupling between cell cycle and cell fate decision may affect the concerted versus sequential dynamical features [11].

In summary, in this work through analyzing scRNA-seq data of CPTs in the context of dynamical systems theory, we identify that many CPTs may share a common concerted mechanism. This conclusion is also supported by an increasing number of studies on various CPT processes reporting existence of intermediate hybrid phenotypes that have co-expression of marker genes of both the initial and final phenotypes such as the partial EMT state [63]. Notice that a cell typically has multiple target phenotypes to choose, functionally the concerted mechanism may allow canalized transition for directing the cells to transit to a specific target phenotype, as visioned by the developmentalist C. H. Waddington [64]. One may perform a more systematic study on the possible classes of CPTs following the procedure developed in this work.

## Glossary

RC: reaction coordinate
PCA: principal component analysis
CPT: cell phenotypic transition
EMT: epithelial-mesenchymal transition
PLSR: partial linear square regression
GRN: gene regulation network
LFDR: local false discover rate
KNN: k-nearest-neighbor

## MATERIALS AND METHODS

### 1. Datasets

The scRNA-seq datasets of dentate gyrus neurogenesis, pancreatic endocrinogenesis, bone marrow hematopoiesis, human glutamatergic neurogenesis , and A549 EMT were obtained from the GEO website with GEO number GSE95753 [28], GSE132188 [54], GSE102698 [46],GSE115813 [24], GSE121861 [48], and, respectively.

### 2 Gene selection and binarization

We focused on genes showing switch-like behavior during the phenotype transition.

1. All genes with highly variable expression levels were selected across the whole dataset [21], according to the following procedure: genes were first grouped into 20 bins based on their mean expression values. Their normalized dispersion was scaled by subtracting the mean of dispersions and dividing standard deviation of dispersions in the corresponding bin.,.
2. The minimum number of shared counts (spliced and un-spliced counts) was set according to the distribution of minimum shared counts (Fig. S10a-c). This value should not be too small to avoid error in the calculation of velocity. Because the datasets have similar distributions of shared counts, we set the minimum number of shared counts as 10.
3. We used a two-step procedure for selecting genes whose expression values were binarized. First, we used Silverman’s bandwidth method to determine whether the expression distribution of a specific gene, estimated with kernel density estimation, is unimodal or multimodal [65]. For the latter case, we calculate the number of modes in the gene expression and find the split values *s_v_*, i.e., the local minima of the kernel density function. Next, we performed K-means clustering to cluster cells into two groups with the gene expressed high and low, respectively. We then used silhouette score to evaluate the clustering quality. A gene was set to be binarizable if: 1) the clustering has a high silhouette score (> 0.65) ; 2) any of the split values is between the centers of the two clusters (*s_v_>c_1_ and s_v_<c_2_*). The reason we combined these two tests is that silhouette score cannot be used to identify the unimodal distribution[66], while the identification of modality through the Silverman’s bandwidth method is sensitive to the fluctuation of data, leading to spurious modes. We applied the procedure to the dentate gyrus neurogenesis, pancreatic endocrinogenesis and human glutamatergic neurogenesis datasets, and identified 678, 470 and 621 genes, respectively.

We followed a different procedure for the hematopoiesis and A549 dataset, since the above two-step procedure only gives 74 and 53 binarized genes. We found that most genes in both datasets have unimodal distributions so fail the *s_v_* test. Instead, we compared the distributions of genes in the 100 cells that are closest to the centers (mean value in the all gene expression space) of initial (HSCs and day 0 sample in bone marrow and A549 dataset separately) and final cell types (monocytes and day 3 sample in bone marrow and A549 dataset separately). If the distributions show significant shift, i.e., the distance between peaks is larger than the summation of the half height peak widths), the gene was selected, and the threshold was calculated by the mean values of the 200 cells (100 cells from the initial cell type and 100 cells from the final cell type). We identified 398 and 437 genes for the hematopoiesis and A549 datasets, respectively.

### 3 Path analysis from single cell RNA velocity analysis

After preprocessing the dataset (Materials and Methods 2), we perform principal component analysis (PCA) for dimension reduction and use scRNA-seq velocity analysis to reconstruct the velocity graph of the whole cell population, which is a transition matrix between all pairs of the cells [31]. The centers (mean values) of the first and last sample populations were calculated, and the distance from each cell to the corresponding center were calculated. For each sample population, the top 100 cells that are closest to the center were selected. We performed single cell trajectory simulation on the transition graph with scVelo [25]. A total of 100 cells were randomly selected from the first sample population. In each simulation, the trajectory started from one of these cells and transited into next cell based on its transition probability in the transition graph. The trajectories that ended in the final sample population were used for the calculation of RCs. For the bone marrow hematopoiesis dataset, the author defined Palantir pseudo-time to characterize the terminal cell states[46]. Here we used it to identify the start and end point of the simulated trajectories because the initial and final cell type have broad distributions. Figure S11 shows some typical gene dynamics along the RCs.

### 4 Procedure for determining RC

We followed a procedure adapted from what used in the finite temperature string method in numerical searching for a one-dimensional RC and non-equilibrium umbrella sampling [32, 34, 67]. The RC was discretized by a set of points that were equally distributed along the RC and divided the *M*-dimensional state space into Voronoi grids, so the value of RC of each cell was assigned by the Voronoi grid that it locates in.

a. Identify the starting and ending points of the reaction path as the means of data points in the state A and state B, respectively. The two points were fixed in the remaining iterations.
b. Construct an initial guess of the reaction path that connects the two ending points in the feature space through linear interpolation. Discretize the path with *N* (*N = 15*) points (called images, and the *k_th_* image denoted as *r*_!_with corresponding coordinate **X**(*r*_k_)) uniformly spaced in arc length.
c. For a given trial RC, divide the multi-dimensional state space by a set of Voronoi polyhedra containing individual images, and calculate the score function, 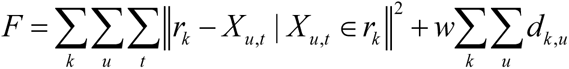, where 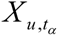 stands for the points on simulated trajectory *u* at step *t_α_* that reside within the *k_th_* polyhedron (containing image point *s_k_*); *d_k,u_* is the distance between image *s_k_* and trajectory *u*, defined as the distance between each image on the path to the closest point on the trajectory, 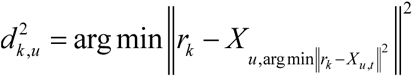; *w* is a parameter that specifies the relative weights between the two terms in the right hand of the expression, here we use 2.
d. Carry out the minimization procedure through an iterative process. For a given trial path defined by the set of image points, we calculate a set of average points using the following equations, 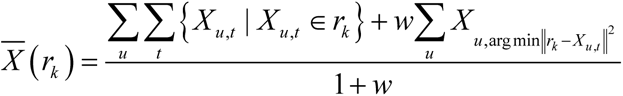. Next we update the continuous reaction path through cubic spline interpolation of the average positions [68], and generated a new set of *N* images {*X* (*r_k_*)} that are uniformly distributed along the new reaction path. We set a smooth factor, *i.e.*, the upper limit of 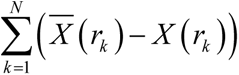, as 1 for calculating the RC.
e. Iterate the whole process in step 3 until there was no further change of Voronoi polyhedron assignments of the data points.

## 5 Network inference

La Manno et al. showed that from scRNA-seq data one can both obtain the single cell expression vectors of the spliced mRNAs {**x***^α^* } , and estimate the instant RNA velocity vectors {**v***^α^* = (*d***x** / *dt*)*^α^*} from reads of spliced and unspliced mRNAs, with α representing the α-th cell [24]. Qiu et al. [30] further developed a procedure of reconstructing the generally nonlinear dynamical equations from the data. We adopted a partial least square regression (PLSR) method to infer the gene regulation networks. We used the velocity vector of each single cell **v** and the level of spliced mRNA ***x*** for inferring the GRN with the PLSR method, ***v*** = ***Fx*** + *error*, where ***F*** is a constant and generally asymmetric matrix describing gene regulation strength. Regression methods are widely used in network inference, and among which the PLSR has several advantages. First, it can be used when the number of features is larger than the number of samples. In the scRNA dataset, the number of genes is often comparable to or larger than the number of cells. Second, it can avoid over-fitting because it uses major components for regression. However, the regulation relation ***F*** obtained from PLSR is typically a dense matrix, while most GRNs are sparse. To generate a sparse network, we further adopt the method of local false discover rate (LFDR) to select those regulation relations that are statistically significant. This procedure ensures the GRN is sparse [35]. Since cells within each Voronoi grid can scatter dispersively in the orthogonal space, we select only cells close to the RC for inferring the ***F*** matrix. That is, among cells within each Voronoi grid, we selected the k-nearest-neighboring (KNN) cells of the corresponding RC point. Such cells from all Voronoi grids collectively form the set for ***F*** matrix inference. We performed the inference using scikit-learn [69] by maximizing the covariance between ***x*** and ***v*** in the PLSR method. The value of components was set to be 2 and data were standardized. In LFDR, the null hypothesis *H* _0_ assumes that *F_i_*_, *j*_ , which is regulation from gene *j* to gene *i*, is 0. An interaction is identified as nonzero when *fdr* (*F_i, j_*) < *q* , where *fdr* (*F_i, j_*) is the false discover rate and *q* is the threshold [70]. The following R package was used for calculation (https://rdrr.io/cran/locfdr/), with a central matching estimation method. Because the *q* value is user controlled, we set the values to make the GRNs in different datasets have similar mean degree numbers. We found the valued doesn’t affect the results in a wide range.

Table S1 shows some of gene regulation pairs inferred with this method. The regulation relations are supported by the literature.

### 6 Calculation of frustration score of GRNs

In this framework, the regulation can be direct such as gene *j* acting as a transcription factor on gene *i*, or indirect that is mediated through molecular species not resolved by the scRNA-seq measurements. While ***F*** is the same for all cells, the expression states of genes within the GRN may vary for different cells. Notice that a prerequisite for gene *j* acting on *i* is that gene *j* is expressed in the cell, otherwise the *j* → *i* edge is treated as non-existent in this specific cell.

Different from frustration in physics, the interactions between genes are directional. We defined a cell-specific effective matrix, *F̄_ij_* = (2*s_i_* −1)*F_ij_* for the definition of frustrated interactions. Then we defined a frustration value for the interaction between a pair of genes (*i*, *j*) as *fs_ij_* = *s _j_* sgn(*F̄_ij_*), assuming a value 1 (not frustrated), 0 (no regulation), and -1 (frustrated), and sgn(x) is the usual sign function,

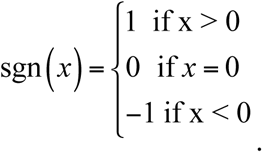

For instance, *fs_ij_* equals 0 if a gene is silent ( *s _j_* = 0), because there is no effective regulation from it. All the regulations from it to other genes will be neither frustrated nor un-frustrated.

### 7 Network Heterogeneity

Gao et al defined network heterogeneity to characterize network topological structures [38]. One defines vectors of outgoing and ingoing weighted degrees of gene connectivity in the network as, *d^out^* = **1***^T^* **F̄** ’, and *d^in^* = **F̄’1** , respectively, with **1** the unit vector **1** = (1, …, 1)*^T^*, and **F̄**’ the **F̄** matrix excluding the diagonal terms. Network heterogeneity measures how homogenously the weighted connections are distributed among the genes, ℋ = *σ^in^σ^out^*⁄〈*d*〉, with *σ^in^* and *σ^out^* the square roots of variances of the elements of *d^in^* and *d^out^*, respectively.

Jacob proposed another definition of measuring network heterogeneity for undirected network. The definition is as follows:

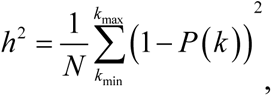

Where *h* is the degree heterogeneity, *N* is the number of nodes in the network, *P* (*k* ) is the probability that a node’s degree is *k* (*P* (*k*) ≠ 0 in this equation). In our calculation, we treat the summation of the in and out edges of one node as its degree value.

### 8 Community Detection

The community detection is performed with Leiden algorithm [40]. In Leiden algorithm, the resolution of community detection is tunable. More communities will be detected when a higher resolution parameter is used. There are communities with only 1 isolated gene (gene that has no interaction with other genes), Because these genes accounts for very small proportion in the whole gene set and they don’t affect the results of inter-community edges, they are not plotted in Fig. 2e, Fig. 3f and Fig. 4e. Gene enrichment analysis of each community is performed with GSEAPY [71, 72]. Table S2-4 show top-ranked inter-community edges.

### 9 Network inference with GRISLI

Another GRN inference method was used to confirm the results. GRISLI treats the GRN inference problem as a sparse regression problem [47]. To solve the equation **v** = **Fx** + *ε*, GRISLI utilizes a stability selection method called TIGRESS (Trustful Inference of Gene Regulation with Stability Selection). TIGRESS is a combination of least angle regression (LARS) with stability selection. A scoring technique was used in stability selection [73]. While GRISLI also proposes a RNA velocity inference method with a spatial-temporal kernel, we still used the velocity values inferred from Dynamo and scVelo to be consistent in comparison.

### 10 Statistical analyses

We randomly selected a subset of cells or genes randomly from the processed dataset and performed the same analyses. The results are not affected by the choice of cells and genes (Fig. S12). We also calculated the number of inter-community edges under different resolution of Leiden algorithm. The resolution value does not affect the results (Fig. S13).

## Code availability

The package *dynamo can be found in* https://dynamo-release.readthedocs.io/en/latest/. And all the scripts are available in https://github.com/opnumten/CPT_GRN_analysis/

## Acknowledgements

We thank Yan Zhang for help discussions on using the *dynamo* package. This work was partially supported by National Cancer Institute (R37 CA232209), and National Institute of Diabetes and Digestive and Kidney Diseases (R01DK119232) to JX, and National Institute of Biomedical Imaging and Bioengineering (T32EB009403) to DP.

## Supplemental figure captions

**Figure S1.**
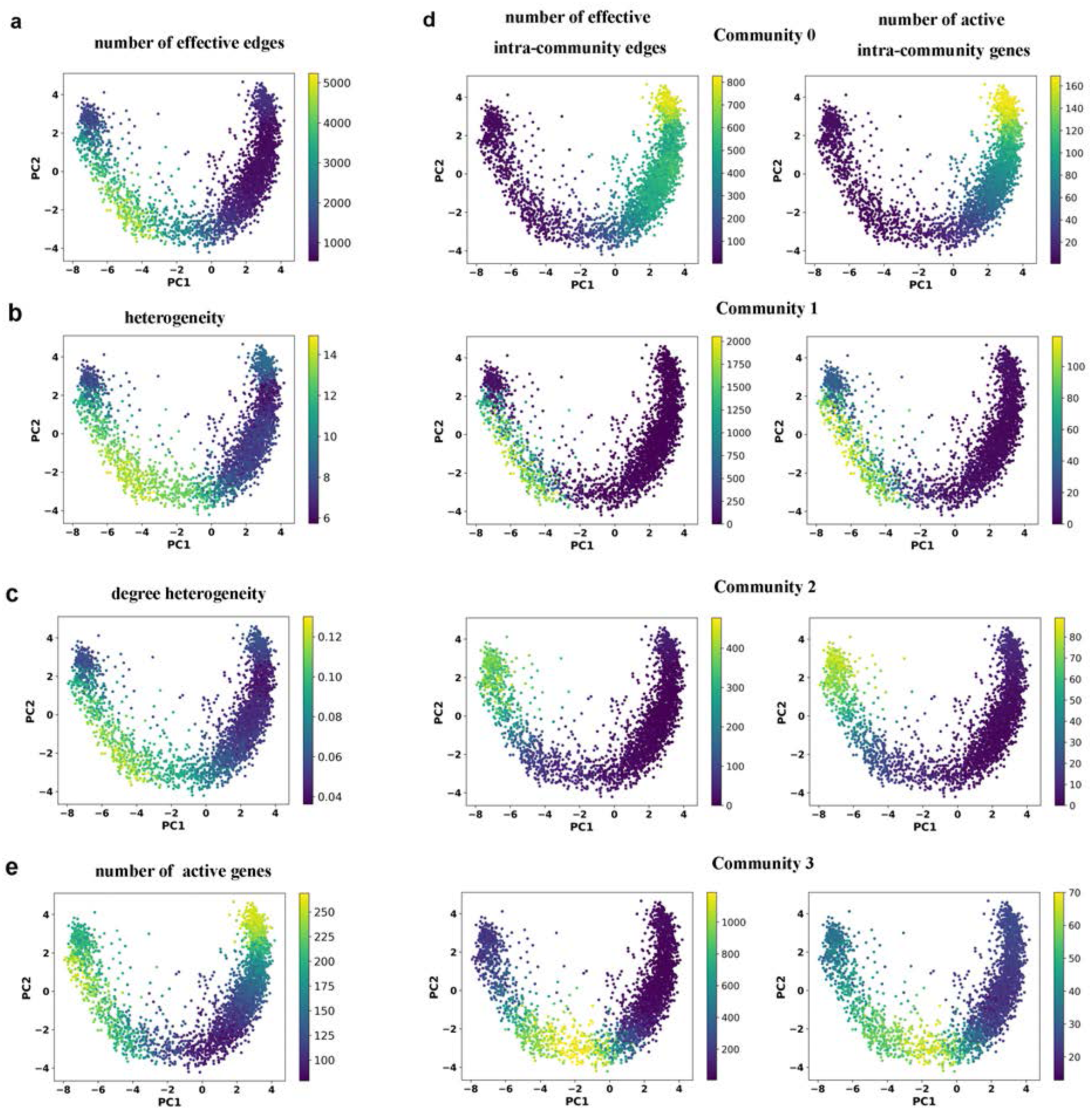
Additional results on transition path analyses of the dentate gyrus neurogenesis dataset. (a) Cell-specific variation of the number of effective regulation edges. Each dot represents a cell. Color represents number effective edges within each cell in the GRN. (b) Cell-specific variation of the network heterogeneity. Each dot represents a cell. Color represents the value of heterogeneity of GRN within each cell. (c) Cell-specific variation of the network degree heterogeneity. Each dot represents a cell. Each dot represents a cell. Color represents the value of degree heterogeneity of GRN within each cell. (d) Left: Cell-specific variation of the number of effective regulation edges inside each community. Each dot represents a cell. Color represents the number of effective edges of the community within each cell. Right: Cell-specific variation of the number of active genes inside each community. Each dot represents a cell. Color represents the number of active genes of the community within each cell. (c) Cell-specific variation of the total number of active genes. Each dot represents a cell. Color represents the total number of active genes within each cell.

**Figure S2.**
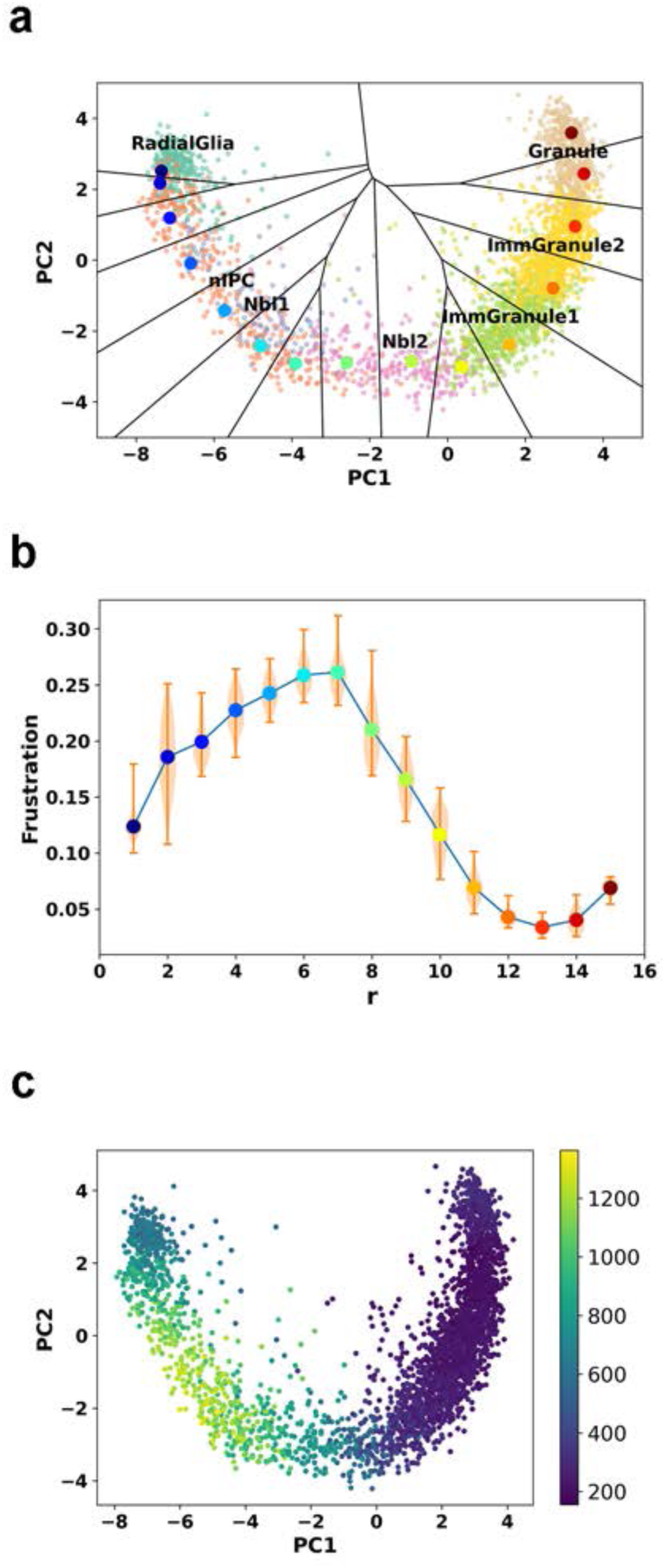
Analysis of the dentate gyrus neurogenesis dataset using the package scVelo. (a) The RC and corresponding Voronoi grids. The large colored dots represent the RC points (starts from blue and ends in red). The small dots represent cells with color as cell type. (b) Frustration score along the RC of dentate gyrus neurogenesis. (c) Variation of effective inter-community regulation edges in the GRN in the processes of dentate gyrus neurogenesis. Color represents the number of effective inter-community regulation edges in individual cell.

**Figure S3.**
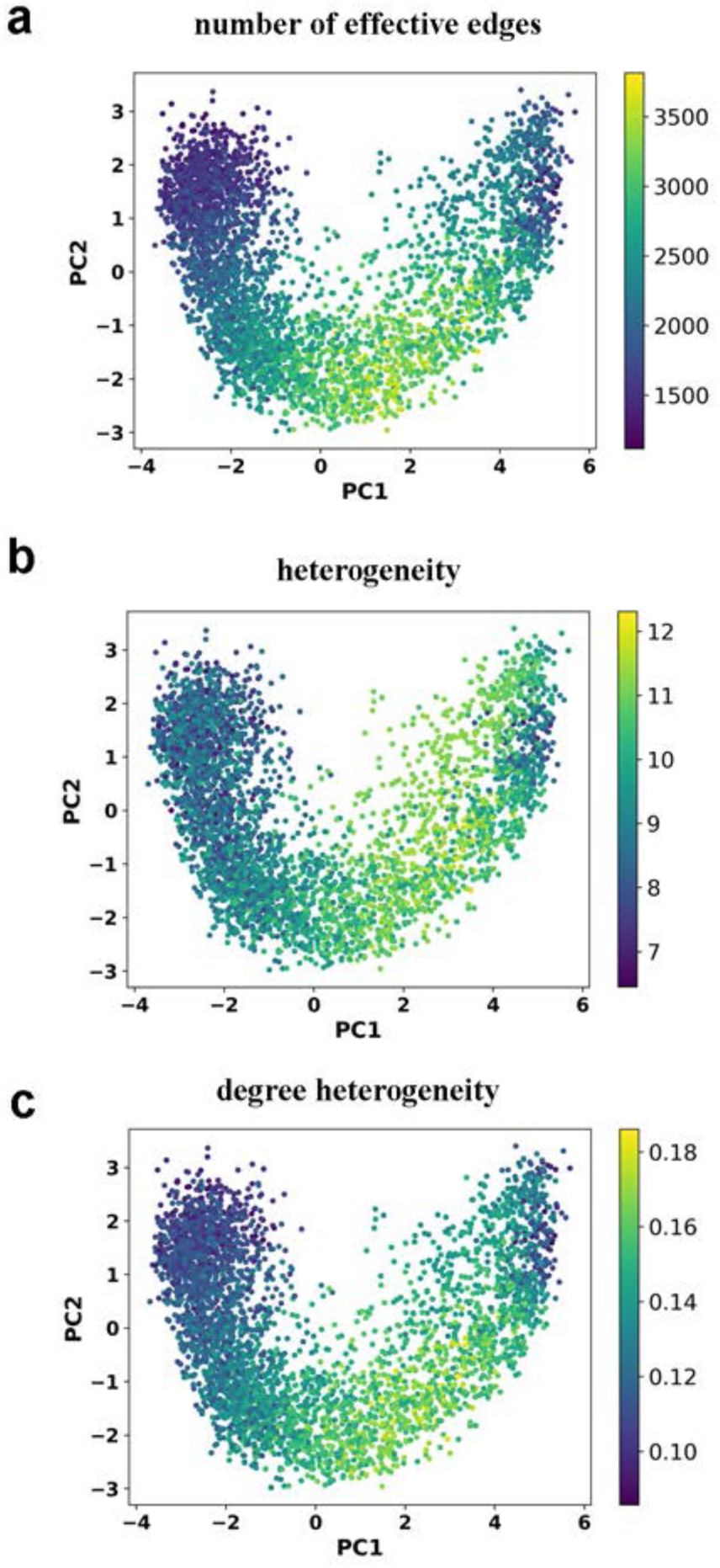
GRN analyses of the hematopoiesis dataset. (a) Cell-specific variation of the number of effective regulation edges. Each dot represents a cell. Color represents number of effective edges in GRN within each cell. (b) Cell-specific variation of heterogeneity. Each dot represents a cell. Color represents value of heterogeneity of GRN within each cell. (c) Cell-specific variation of degree heterogeneity. Each dot represents a cell. Color represents value of degree heterogeneity of GRN within each cell.

**Figure S4.**
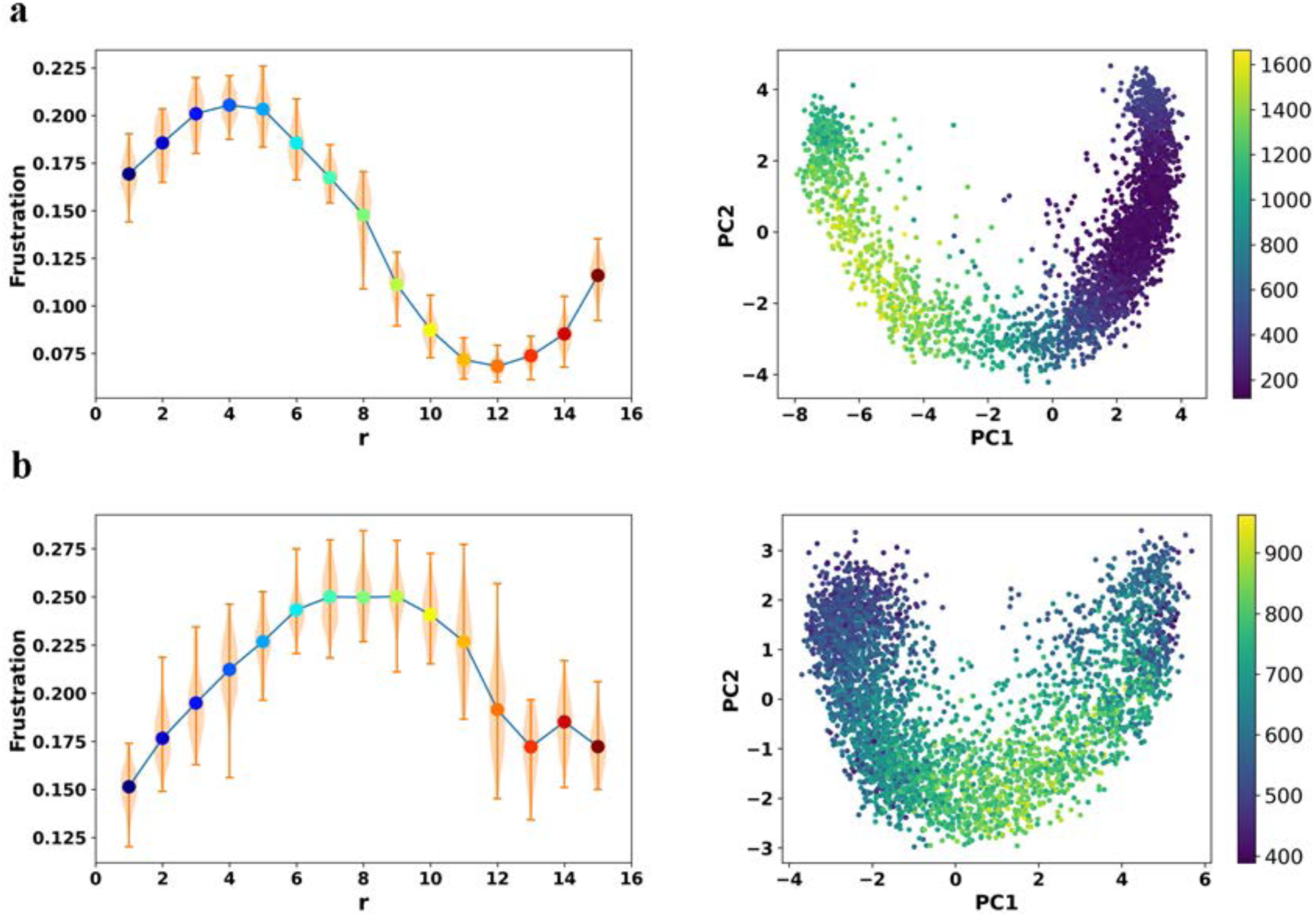
Analyses of datasets of dentate gyrus neurogenesis (a) and hematopoiesis (b) based on GRN inferred with GRISLI. (a) Left: Frustration score along the RC of dentate gyrus neurogenesis; Right: cell-specific variation of the number of inter-community edges. Each dot represents a cell and color represents the number of inter-community edges in GRN within each cell. (b) Same as in panel (a), except for hematopoiesis.

**Figure S5.**
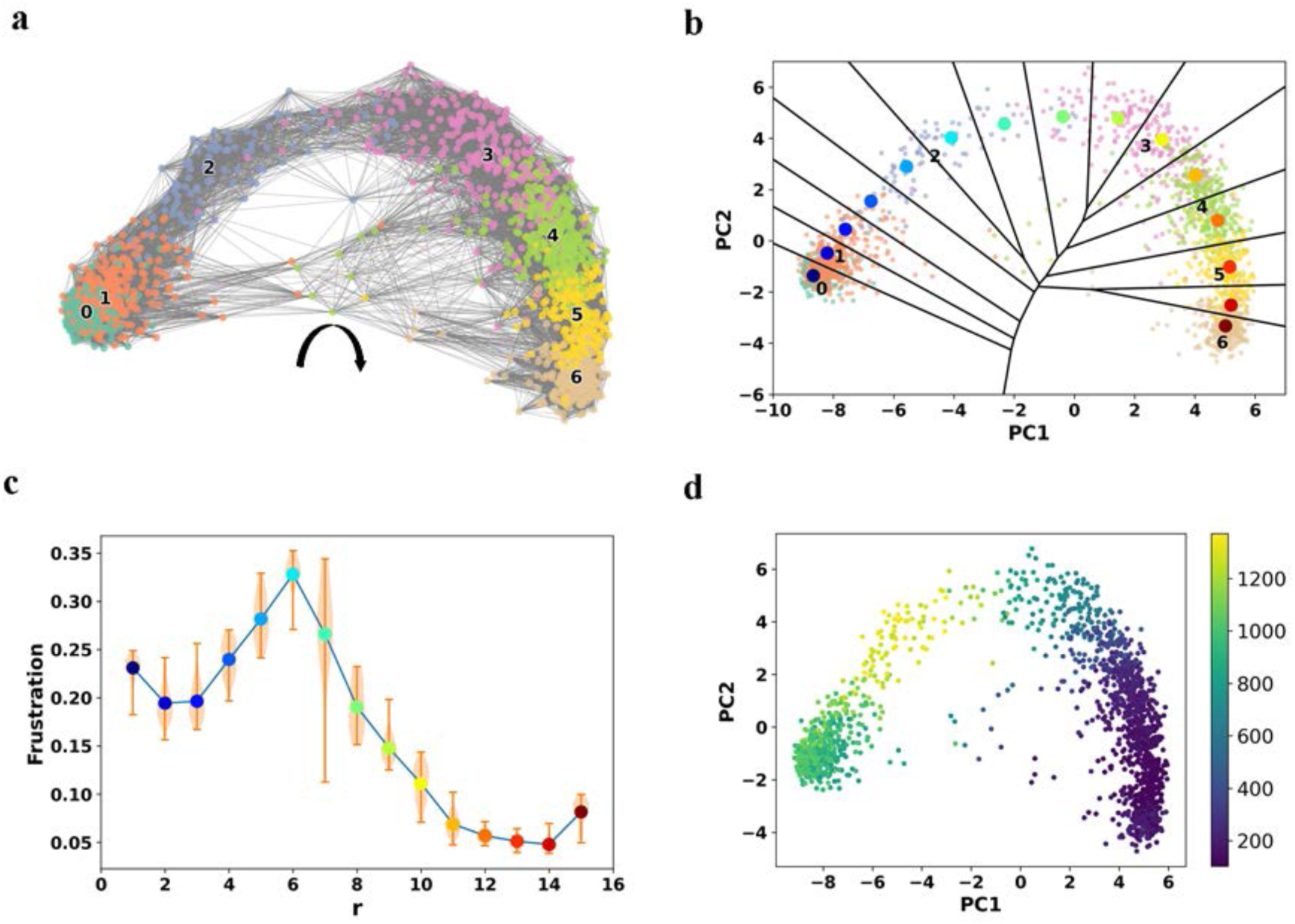
Analyses on human glutamatergic neurogenesis. (a) Transition graph of development of human glutamatergic neuronal lineage based on RNA velocity. Arrow represents the direction of development. 0-1,2-3, 4 and 5-6 correspond to radial glia, neuroblast, immature neuron and neuron separately. (b) The RC and corresponding Voronoi grids. The large colored dots represent the RC points (start from blue and end in red). The small dots represent cells with color indicating cell type. (c) Frustration score along the RC in development of human glutamatergic neuronal lineage. (d) Cell-specific variation of effective intercommunity regulation in development of human glutamatergic neuronal lineage. Each dot is a cell. Color represents the number of effective intercommunity edges within each cell in the GRN.

**Figure S6.**
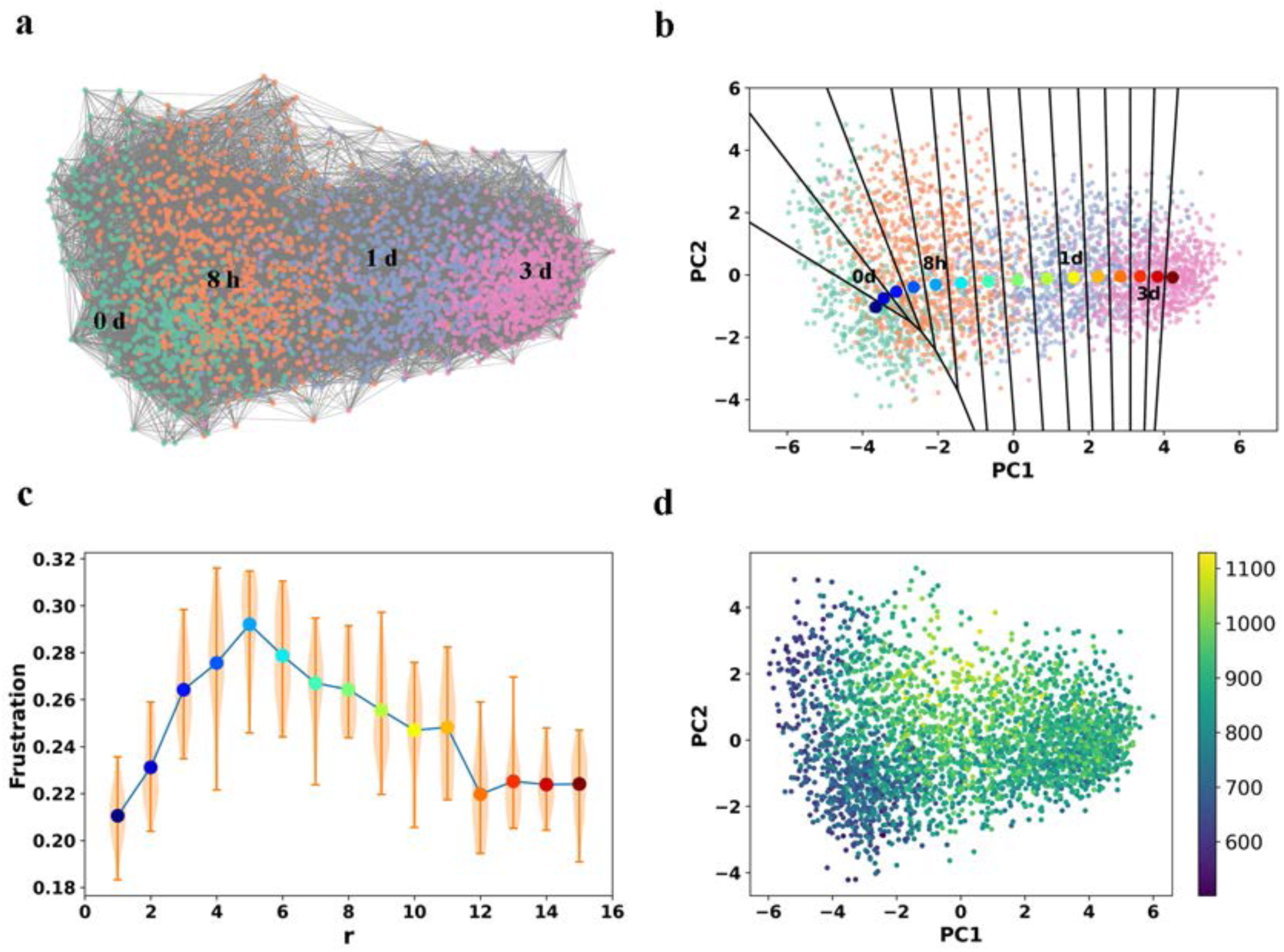
Analyses on EMT of A549 cells treated with TGF-β. (a) Transition graph of EMT. Each dot is a single cell. Color represents the duration of TGF-β treatment (0 day, 8 hour, 1 day and 3 day). (b) The RC and corresponding Voronoi grids. The large colored dots represent the RC points (start from blue and end in red). The small dots represent cells with color as cell type. (c) Frustration score along the RC. (d) Cell-specific variation of effective intercommunity regulation in EMT. Each dot is a cell. Color represents the number of effective intercommunity edges within each cell in the GRN.

**Figure S7.**
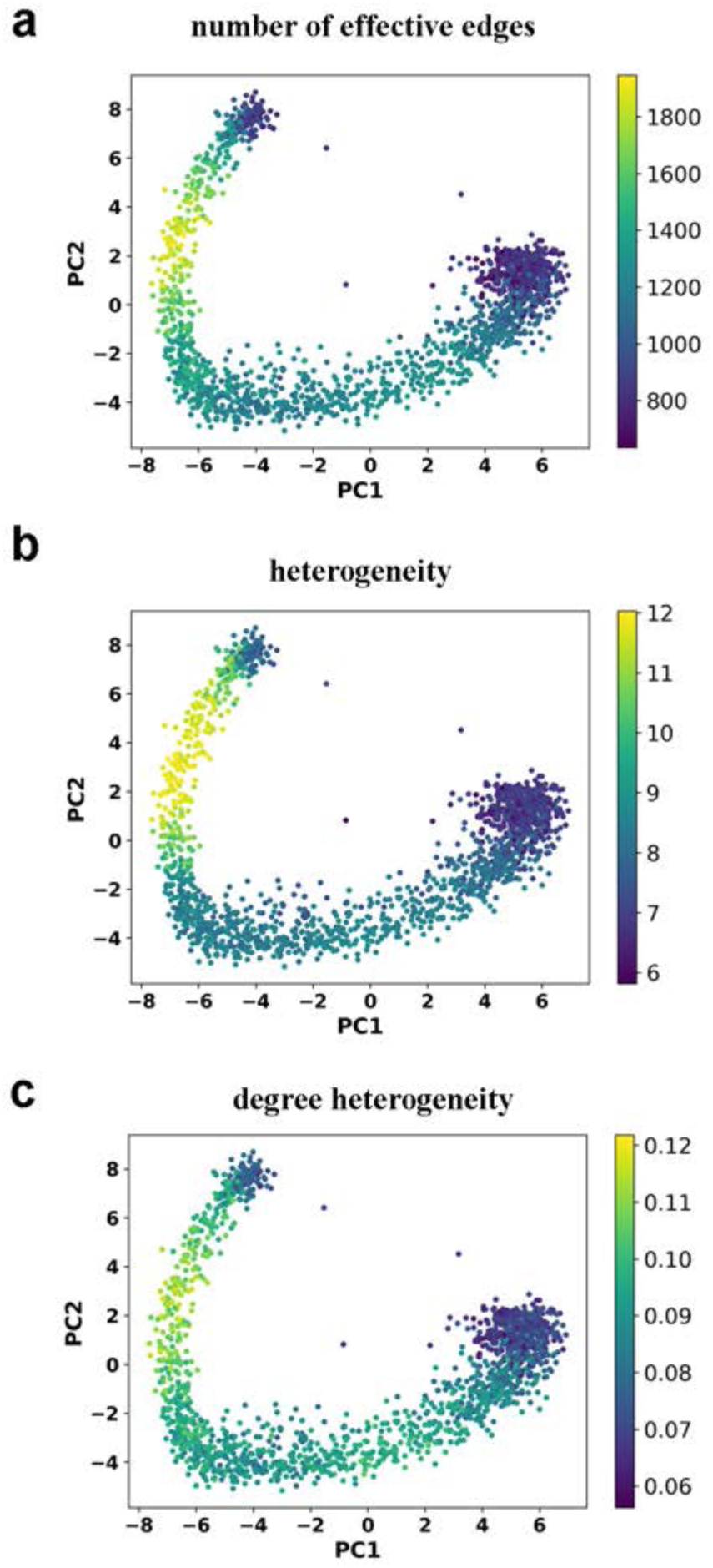
GRN analyses of pancreatic endocrinogenesis. (a) Cell-specific variation of the number of effective regulation edges. Each dot represents a cell. Color represents number of effective edges in GRN within each cell (b) Cell-specific variation of heterogeneity. Each dot represents a cell. Color represents value of heterogeneity of GRN within each cell. (c) Cell-specific variation of degree heterogeneity. Each dot represents a cell. Color represents value of degree heterogeneity of GRN within each cell.

**Figure S8.**
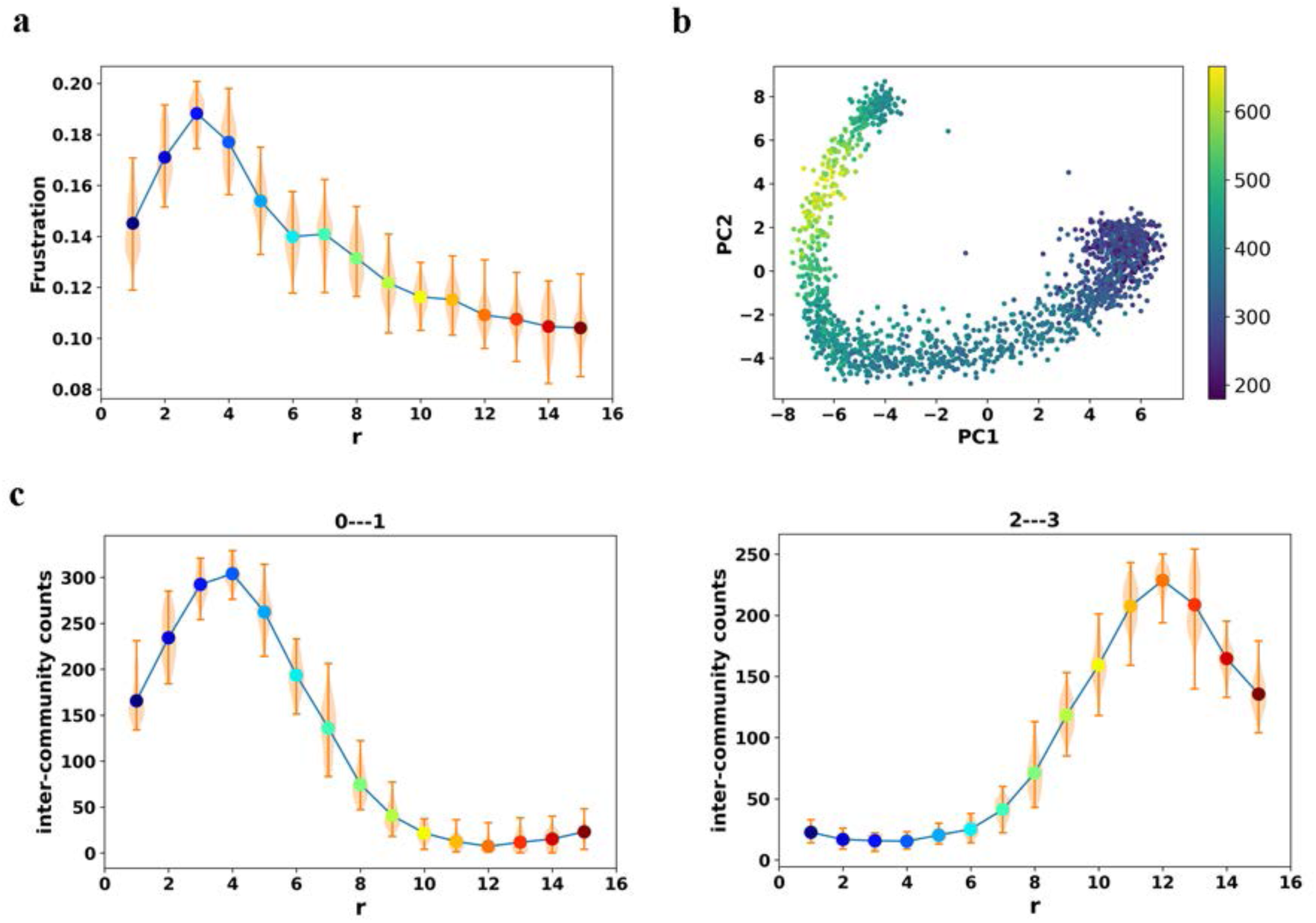
Analyses of pancreatic endocrinogenesis based on GRN inferred with GRISLI. (a) Frustration score along the RC of pancreatic endocrinogenesis. (b) Cell-specific variation of the number of inter-community edges. Each dot represents a cell and color represents the number of inter-community edges in GRN within each cell. (c) Number of effective intercommunity edges between community 0 and 1 (left) and that between community 2 and 3 (right) along the RC during pancreatic endocrinogenesis.

**Figure S9.**
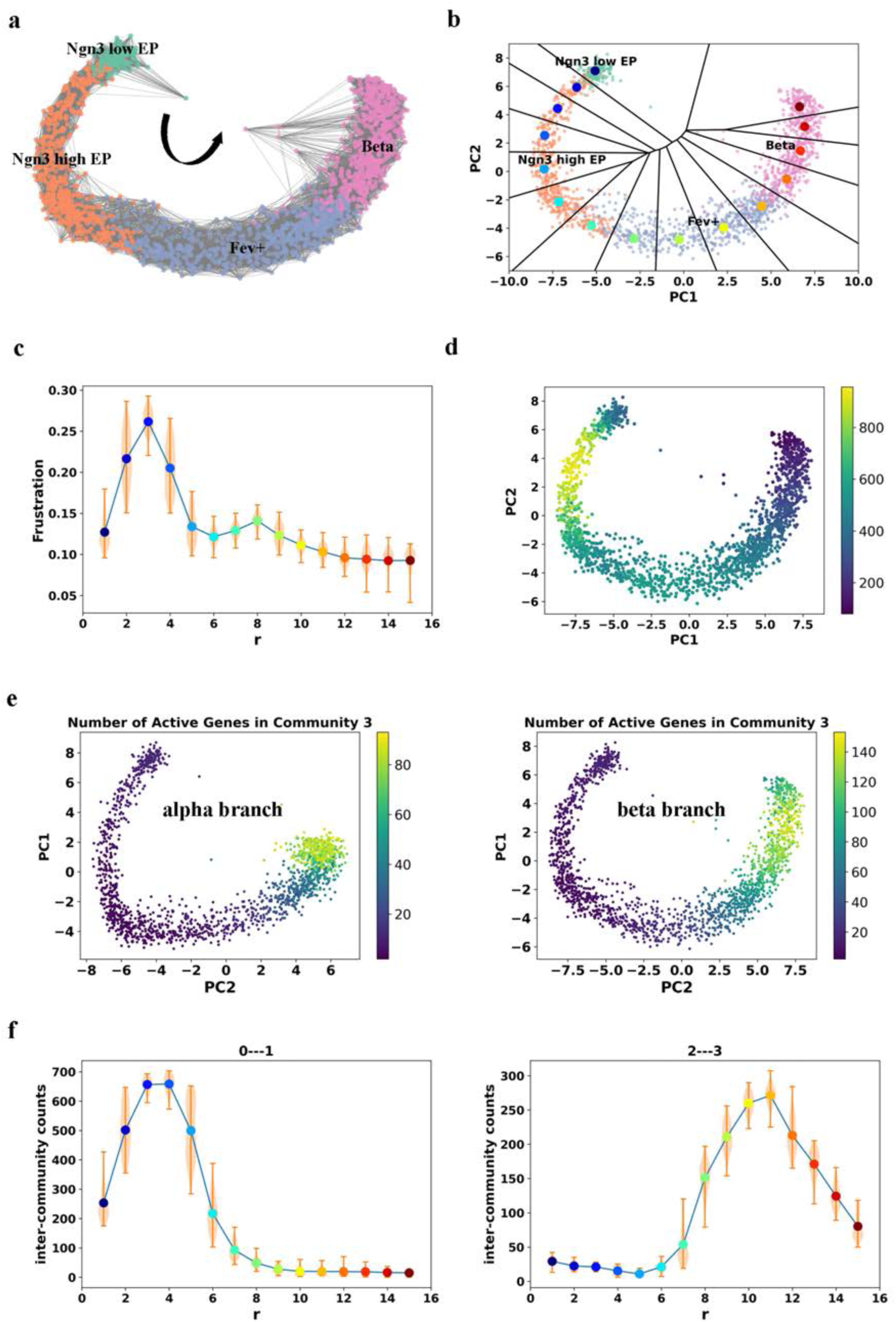
Analyses on the branch of glucagon producing β-cells in pancreatic endocrinogenesis. (a) Transition graph based on RNA velocity. (b) The RC and corresponding Voronoi grids. The large colored dots represent the RC points (start from blue and end in red). The small dots represent cells with color as cell type. (c) Frustration score along the RC. (d) Cell-specific variation of effective intercommunity regulation. Each dot represents a cell. Color represents the number of effective intercommunity edges within each cell in the GRN. (e) Cell-specific variation of the number of active genes inside community 3 in α (left) and β branches (right). Each dot represents a cell. Color represents the number of active genes of the community within each cell. (f) Number of effective intercommunity edges between community 0 and 1 (left) and that between community 2 and 3 (right) along the RC during pancreatic endocrinogenesis in the β branch.

**Figure S10.**
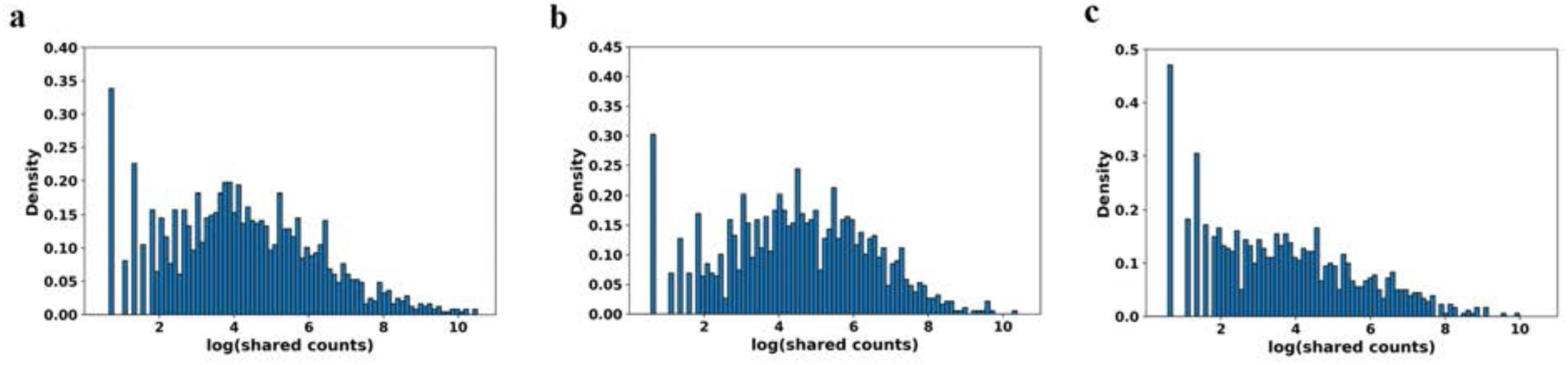
Shared counts distribution of the datasets. (a) Dentate gyrus neurogenesis; (b) Bone marrow hematopoiesis; (c) Pancreatic endocrinogenesis.

**Figure S11.**
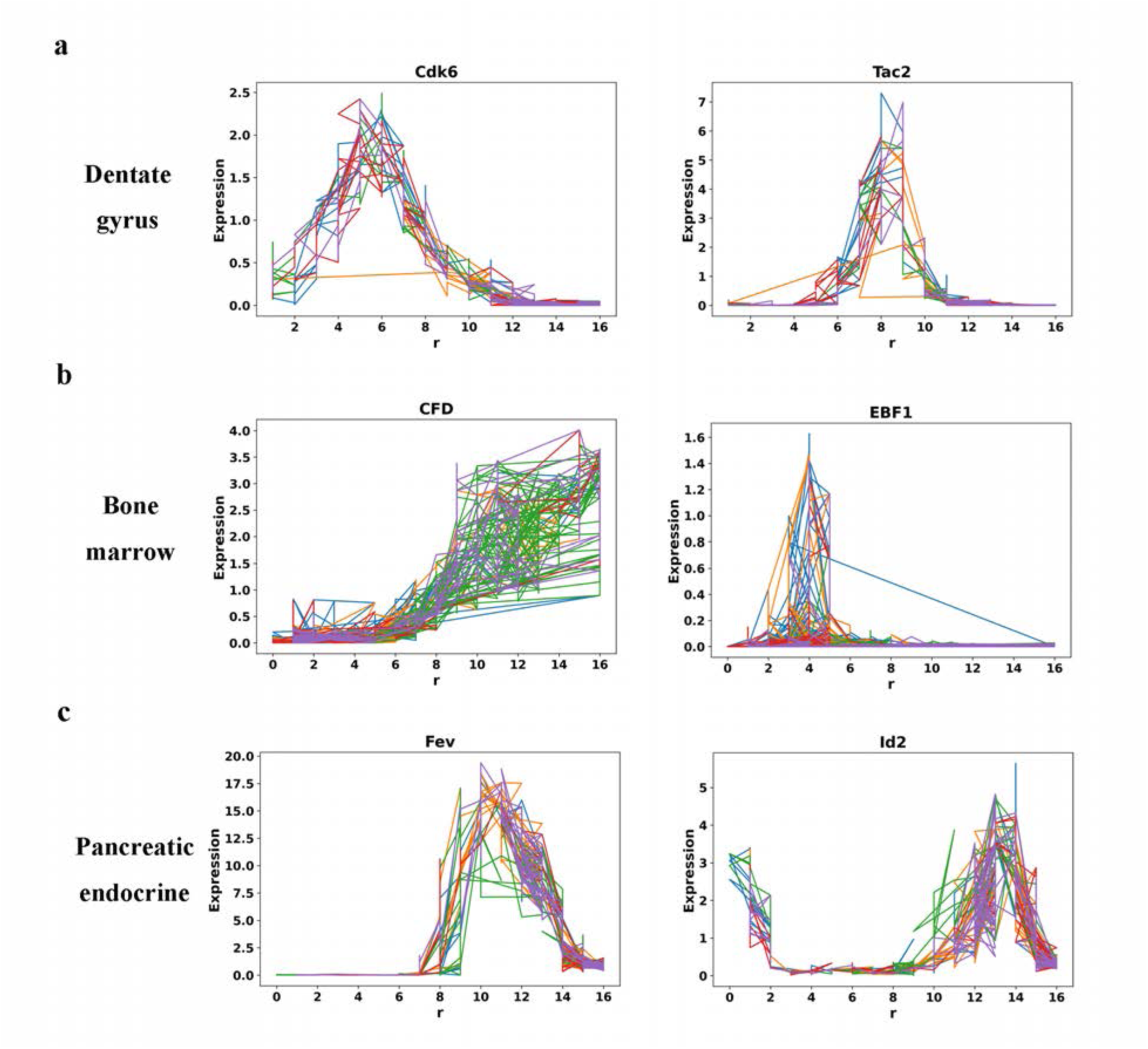
Typical trajectories of high variance genes versus RCs of dentate gyrus neurogenesis (a), bone marrow hematopoiesis (b) and pancreatic endocrinogenesis (c).

**Figure S12.**
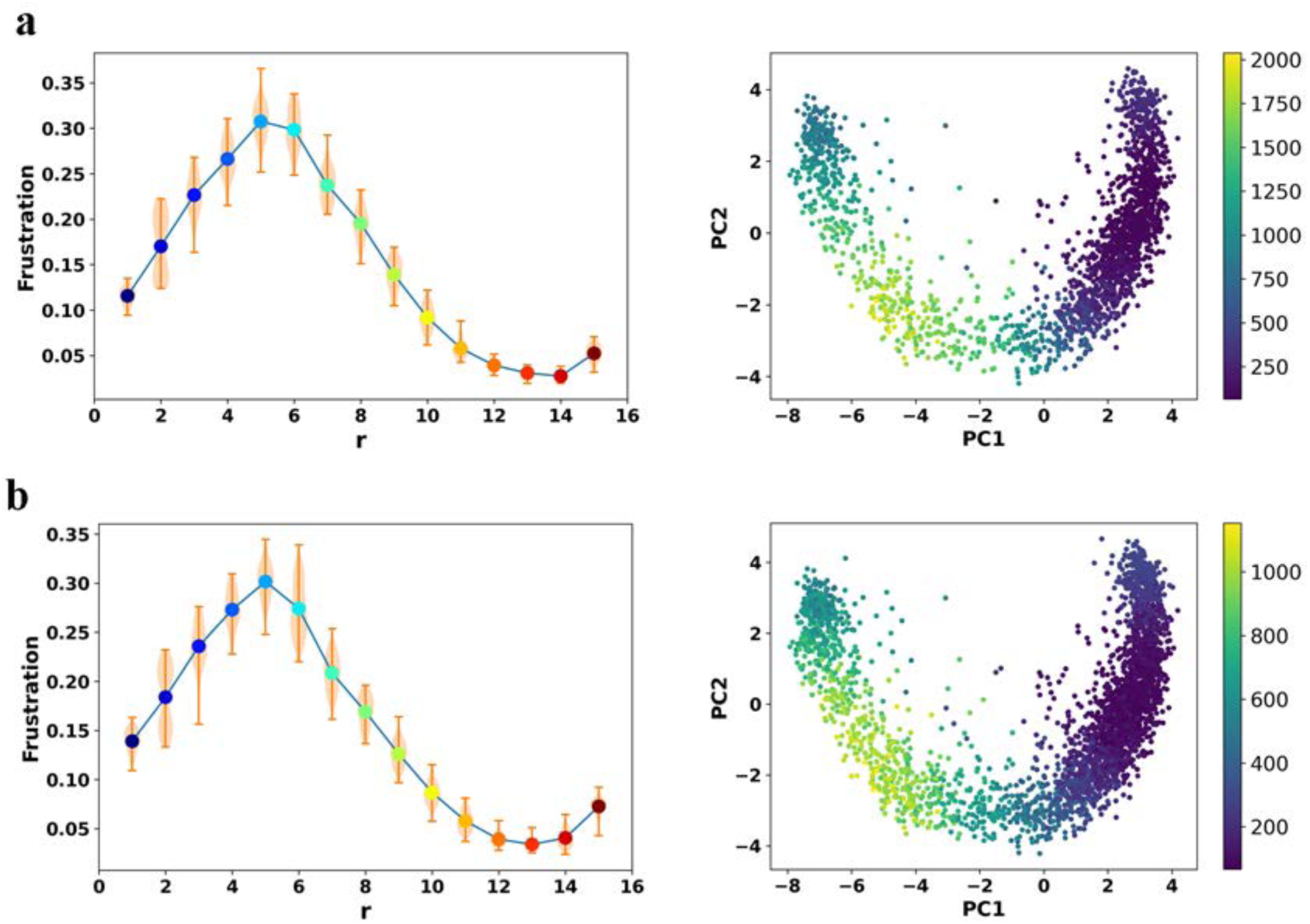
Statistical analyses of dentate gyrus neurogenesis. Each dot represents a cell and color represents the number of inter-community edges. (a) Frustration score along the RCs (left) and cell-specific variation of the number of inter-community edges (right) of a randomly selected sub-population of 2000 cells (from a total of 3184 cells). (b) Frustration score along the RCs (left) and cell-specific variation of the number of inter-community edges) (right) of cells on the space of 400 randomly selected genes (from a total of 678 genes).

**Figure S13.**
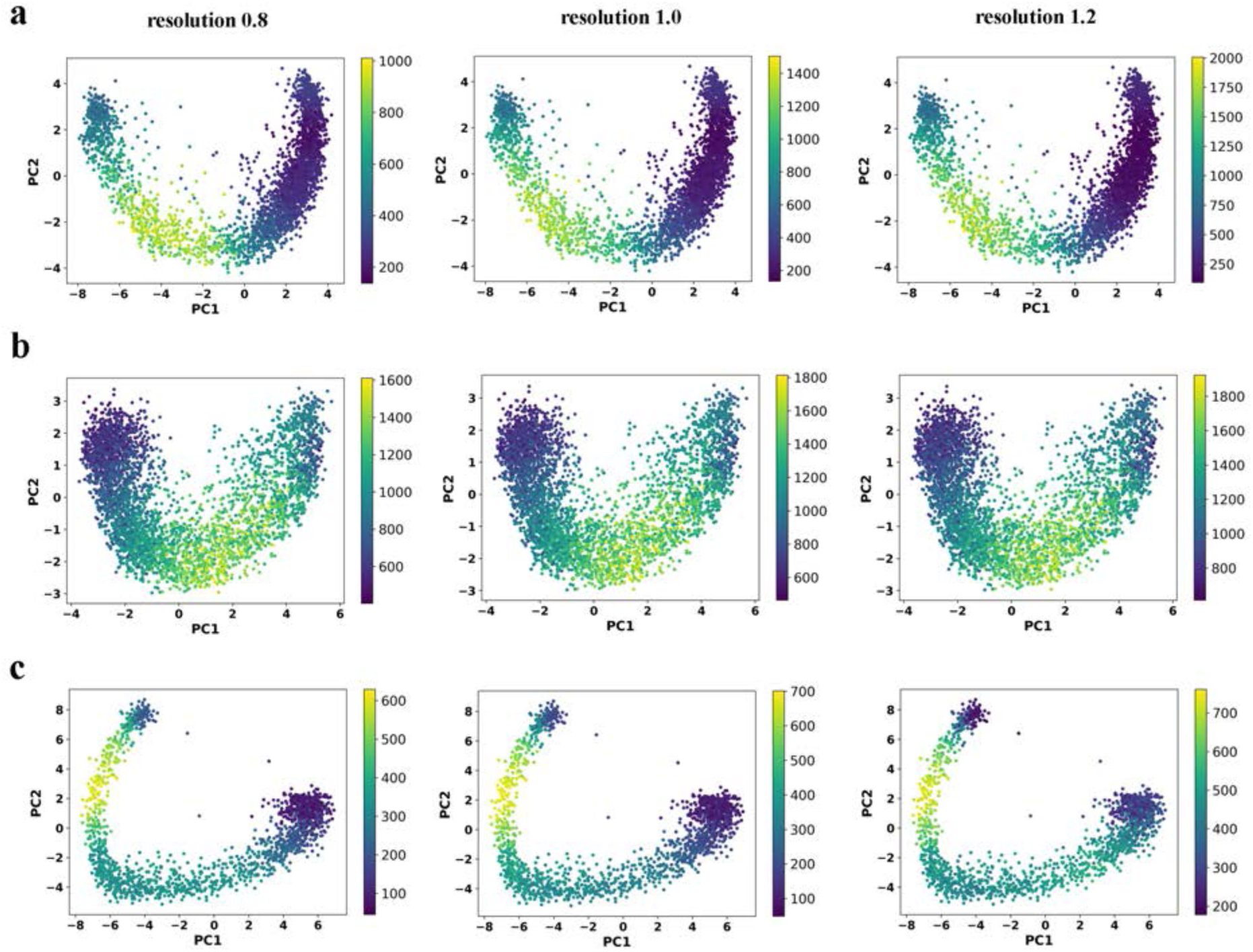
Cell-specific variation of the number of effective inter-community edges between communities calculated with different resolution parameter values for dentate gyrus neurogenesis (a), bone marrow hematopoiesis (b), and pancreatic endocrinogenesis (c). Each dot represents a cell and the color represents the number of inter-community edges.

***Movie S1 Evolution of the number of effective intercommunity edges along the RC during pancreatic endocrinogenesis. Each node represents a gene. Color of the node represents index of community. Arrow represents direction of regulation. r is the index in the RC*.**

**Table S1.**
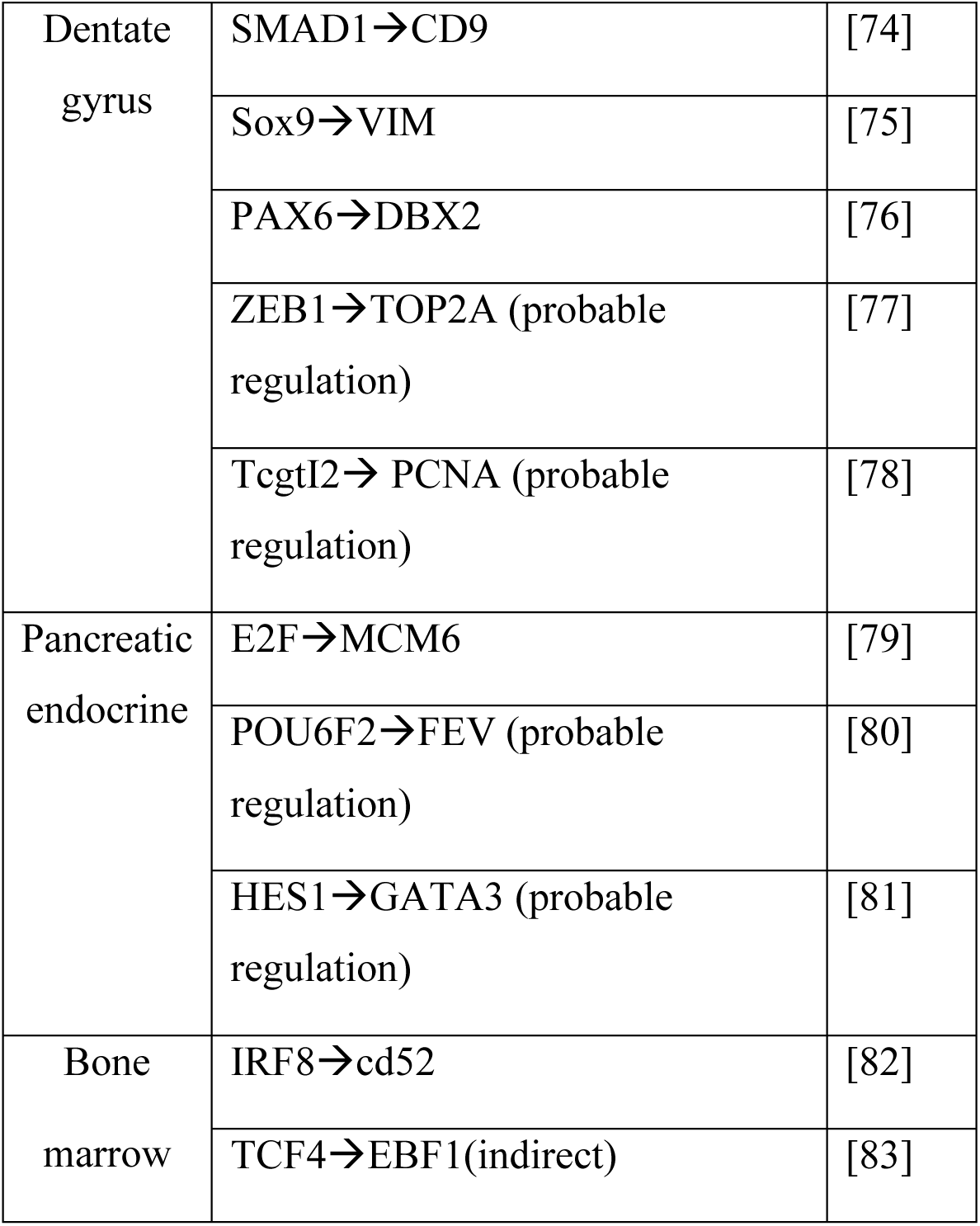
Example strong gene regulation interactions inferred with the PLSR method. The third column are the references that support the regulation in the same row. Probable regulation means there is a possible interaction between the two genes suggested by corresponding literature. Indirect means the regulation is probably indirect as suggested by the literature.

**Table S2.**
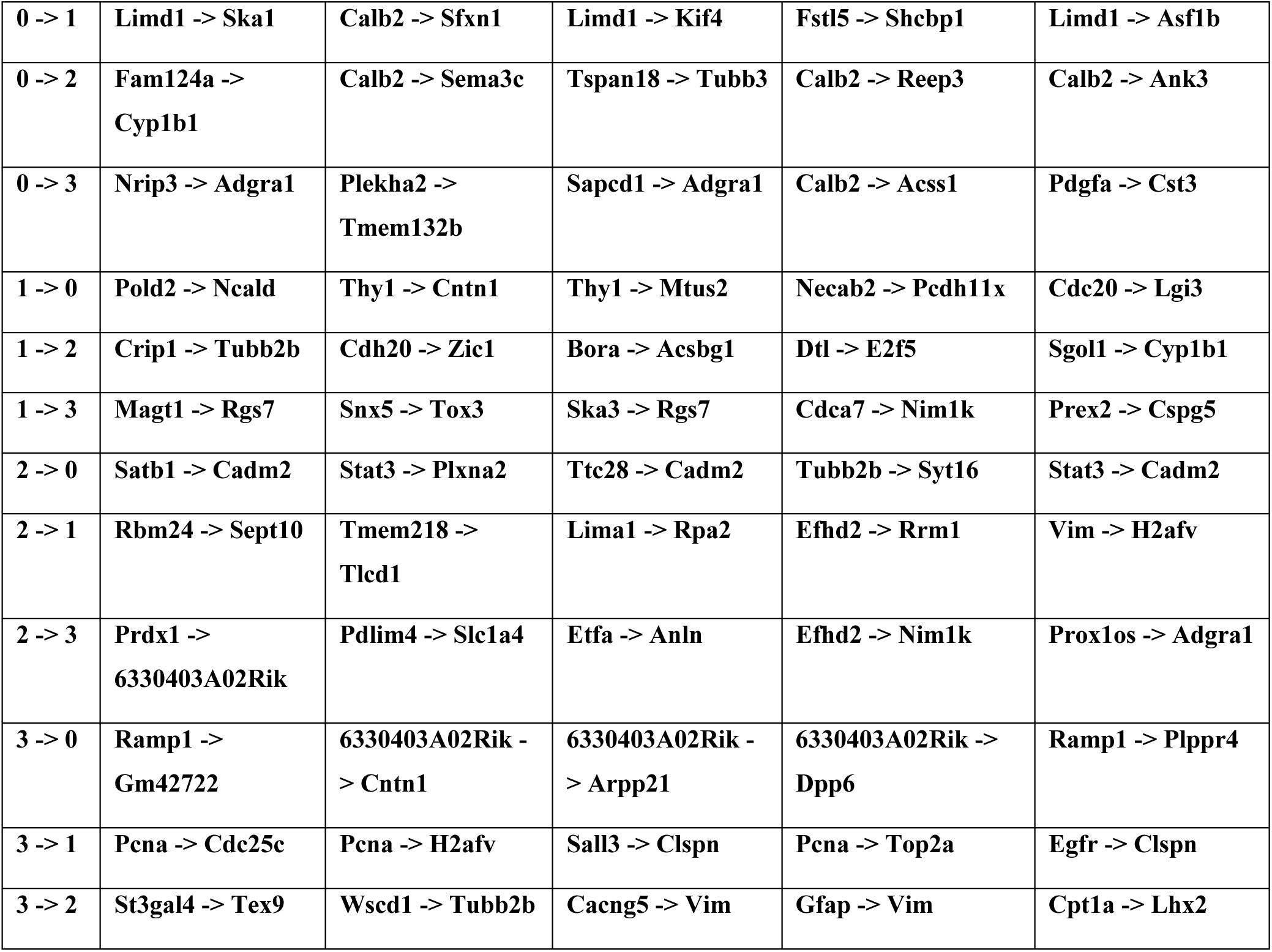
Inter-community interactions with highest weights in dentate gyrus dataset. Gene enrichment analysis reveals that community 0 includes genes related to neural functions such as axon guidance and glutamatergic synapse. Community 1 includes genes related to cell cycle and DNA repair.

**Table S3.** Inter-community interactions with highest weights in hematopoiesis dataset. Gene enrichment analysis reveals that community 0 includes genes related to cardiac functions. Community 2 includes genes related to various signaling pathways such as calcium.

|  |  |  |  |  |  |
| --- | --- | --- | --- | --- | --- |
| 0 -> 1 | ACVR2A -><br>GCSAML | FAM161A -><br>RP11-244M2.1 | CECR1 -> HPGD | JCHAIN -><br>SMAGP | LGMN -><br>HPGD |
| 0 -> 2 | NFIL3 -> PLAC8 | KIF14 -><br>ARHGAP11B | KIF14 -> SCLT1 | HLA-DRB5 -><br>BACH2 | DEPDC1 -><br>CLEC2B |
| 1 -> 0 | AC004791.2 -><br>ZBTB20 | CTSG -> PRR7 | AP4B1 -><br>FCER1A | RASGRP3 -><br>ANGPT2 | CA2 -> DYX1C1 |
| 1 -> 2 | CTSG -> CFD | PPIB -> LYST | RASGRP3 -><br>HLA-DMB | RP11-84C10.2 -><br>SCPEP1 | PRTN3 -><br>RGS18 |
| 2 -> 0 | POU2F2 -><br>LINC01122 | SCLT1 -><br>ZBTB20 | RNH1 -> MDK | SCPEP1 -> MDK | SGTB -><br>MMRN1 |
| 2 -> 1 | LYZ -> PPP1R9A | RNH1 -> FGF13 | LYZ -> AZU1 | ECT2 -> ZNRF3 | SERPINB8 -><br>HPGD |

**Table S4.**
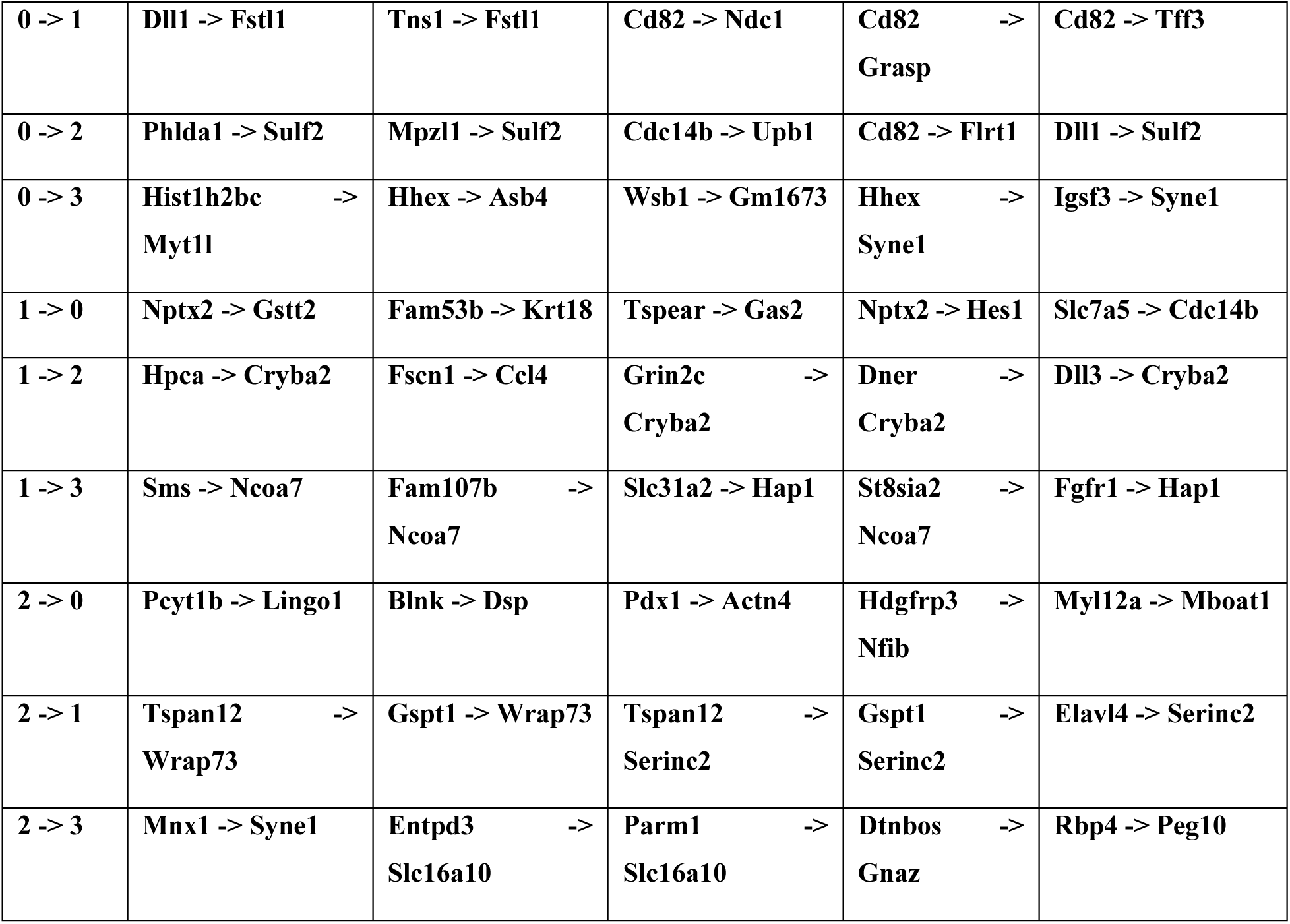
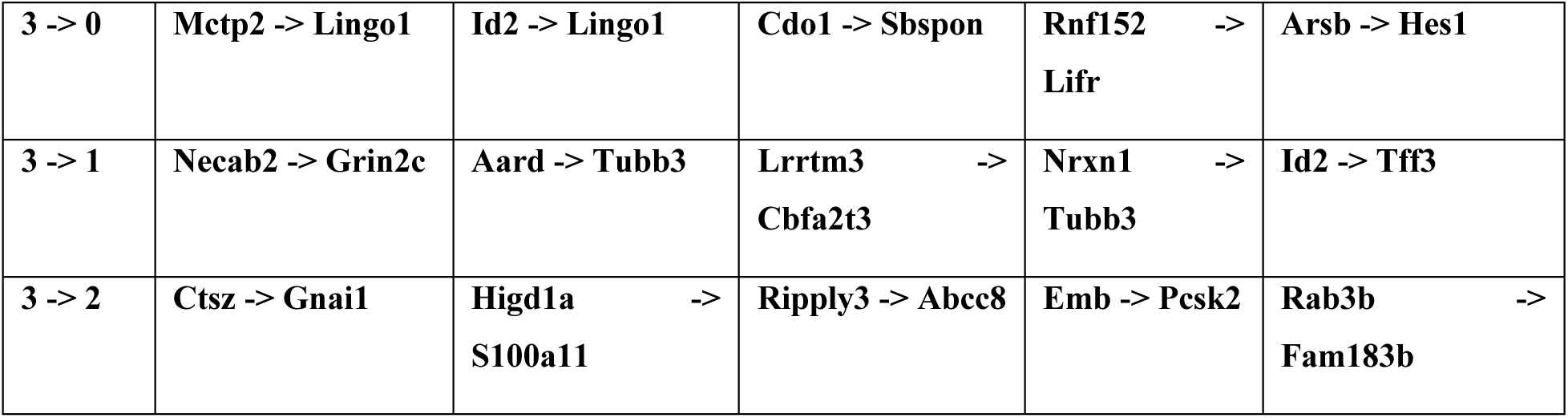
Inter-community interactions with highest weights in pancreatic endocrine dataset. Gene enrichment analysis reveals that community 2 includes genes related to pancreatic functions such as insulin secretion.

